# Neuronal upregulation of Prospero protein is driven by alternative mRNA polyadenylation and Syncrip-mediated mRNA stabilisation

**DOI:** 10.1101/135848

**Authors:** Tamsin J. Samuels, Yoav Arava, Aino I. Järvelin, Francesca Robertson, Jeffrey Y. Lee, Lu Yang, Ching-Po Yang, Tzumin Lee, David Ish-Horowicz, Ilan Davis

**Affiliations:** Department of Biochemistry, The University of Oxford, UK; Technion, Haifa, Israel; Janelia Research Campus, Virginia, USA; MRC Laboratory for Molecular Cell Biology, University College, London, UK

## Abstract

During *Drosophila* and vertebrate brain development, the conserved transcription factor Prospero/Prox1 is an important regulator of the transition between proliferation and differentiation. Prospero level is low in neural stem cells and their immediate progeny, but is upregulated in larval neurons and it is unknown how this process is controlled. Here, we use single molecule fluorescent *in situ* hybridisation to show that larval neurons selectively transcribe a long *prospero* mRNA isoform containing a 15 kb 3’ untranslated region, which is bound in the brain by the conserved RNA-binding protein Syncrip/hnRNPQ. Syncrip binding increases the mRNA stability of the long *prospero* isoform, which allows an upregulation of Prospero protein production. Our findings highlight a regulatory strategy involving alternative polyadenylation followed by differential post-transcriptional regulation.

## INTRODUCTION

The central nervous system (CNS) consists of a huge diversity and number of neurons that originate from a limited population of neural stem cells (NSCs) (Kelava and Lancaster, 2016), also known as neuroblasts (NBs) in *Drosophila*. To produce a normal CNS, NBs must divide the correct number of times and their progeny must undergo a precisely regulated programme of differentiation. Many factors and mechanisms regulating these processes have been studied in great detail, and are extensively conserved between vertebrates and *Drosophila* (Homem and Knoblich, 2012). The emphasis in the field has been on identifying key transcription factors and their downstream transcriptional effects. However post-transcriptional regulation, which can modulate protein expression with enhanced spatial and temporal precision, has been less well characterised.

In *Drosophila* embryos and larvae, NBs undergo repeated asymmetric divisions, maintaining a single large cell that retains its stem cell properties. Each division of a type I NB also produces a ganglion mother cell (GMCs), which divides only once to generate a pair of neurons (Homem and Knoblich, 2012; Knoblich, 2008). Type I NB lineage differentiation is regulated by the conserved homeodomain-containing transcription factor, Prospero (Pros)/Prox1 (Bayraktar et al., 2010). Pros activates the expression of genes required for the differentiation of type I NB progeny and suppresses the expression of genes that promote stem cell-like properties (Bello et al., 2006; Betschinger et al., 2006; Choksi et al., 2006; Doe et al., 1991; Lai and Doe, 2014; Lee et al., 2006; Matsuzaki et al., 1992; Vaessin et al., 1991). *pros* mRNA and Pros protein are expressed in the NBs, but are excluded from the nucleus and segregated asymmetrically into the GMC during NB division (Hirata et al., 1995; Kitajima et al., 2010; Knoblich et al., 1995; Spana and Doe, 1995). In this way the sub-cellular localisation of Pros allows the stem cell-like properties of NBs to be maintained, while ensuring their GMC progeny differentiate correctly. In embryonic type I NBs, Pros is expressed in GMCs and new born neurons but is quickly switched off as neurons mature (Srinivasan et al., 1992). In contrast, Pros expression is upregulated in larval neurons and is required to maintain neuronal identity (Carney et al., 2013), but the mechanism controlling this upregulation is not known.

Here, we examine the mechanism of upregulation of Pros expression in larval neurons using single molecule fluorescent *in situ* hybridisation (smFISH) and immunofluorescence (IF) in whole-mount brains (Yang et al., 2017b). We observe hugely increased *pros* mRNA expression in neurons, correlated with upregulated Pros protein. This highly expressed neuronal *pros* mRNA includes an unusually long 15 kb 3’ untranslated region (UTR) (*pros^long^*). *pros^long^* is not produced in embryos or in larval NBs, but is switched on in larval neurons. We show that *pros^long^* is stabilised by binding to the conserved RNA-Binding Protein (RBP), Syncrip (Syp)/hnRNPQ (Kuchler et al., 2014; Liu et al., 2015; McDermott et al., 2014). We find that the *pros* UTR extension is required for the neuronal upregulation of Pros protein. Our observations highlight a novel example of alternative polyadenylation followed by differential post-transcriptional regulation by mRNA stability, that controls the level of protein expression in distinct cell types.

## RESULTS

### Upregulation of Pros protein in neurons is achieved through cell type-specific stabilisation of *pros* mRNA

Pros protein is expressed at low levels in larval type I NBs and GMCs, where it promotes GMC differentiation, and is then upregulated in larval neurons (Carney et al., 2013; Choksi et al., 2006). To determine whether the upregulated Pros expression in neurons is driven by an increase in *pros* mRNA levels or by increased Pros translation, we used smFISH in whole-mount *Drosophila* larval brains at the wandering third instar stage (Yang et al., 2017b). We carried out four colour imaging with smFISH against *pros* exon (Figure 1A), anti-Pros antibody, anti-Elav antibody (marking differentiated neurons) and DAPI (marking DNA in all cells) (Figure 1B). NBs were easily identified by their large cell size. As previously shown (Carney et al., 2013; Choksi et al., 2006), we found that Pros expression is low in NBs and GMCs (outlined in white and pink respectively) and is upregulated in Elav+, post-mitotic neurons (Figure 1B). *pros* exon smFISH signal is also upregulated in Elav^+^ cells, and correlates with the increased Pros protein expression (Figure 1B). We conclude that increased *pros* transcript number is responsible for the increased Pros protein expression in the neurons compared to the NB.

**Figure 1:**
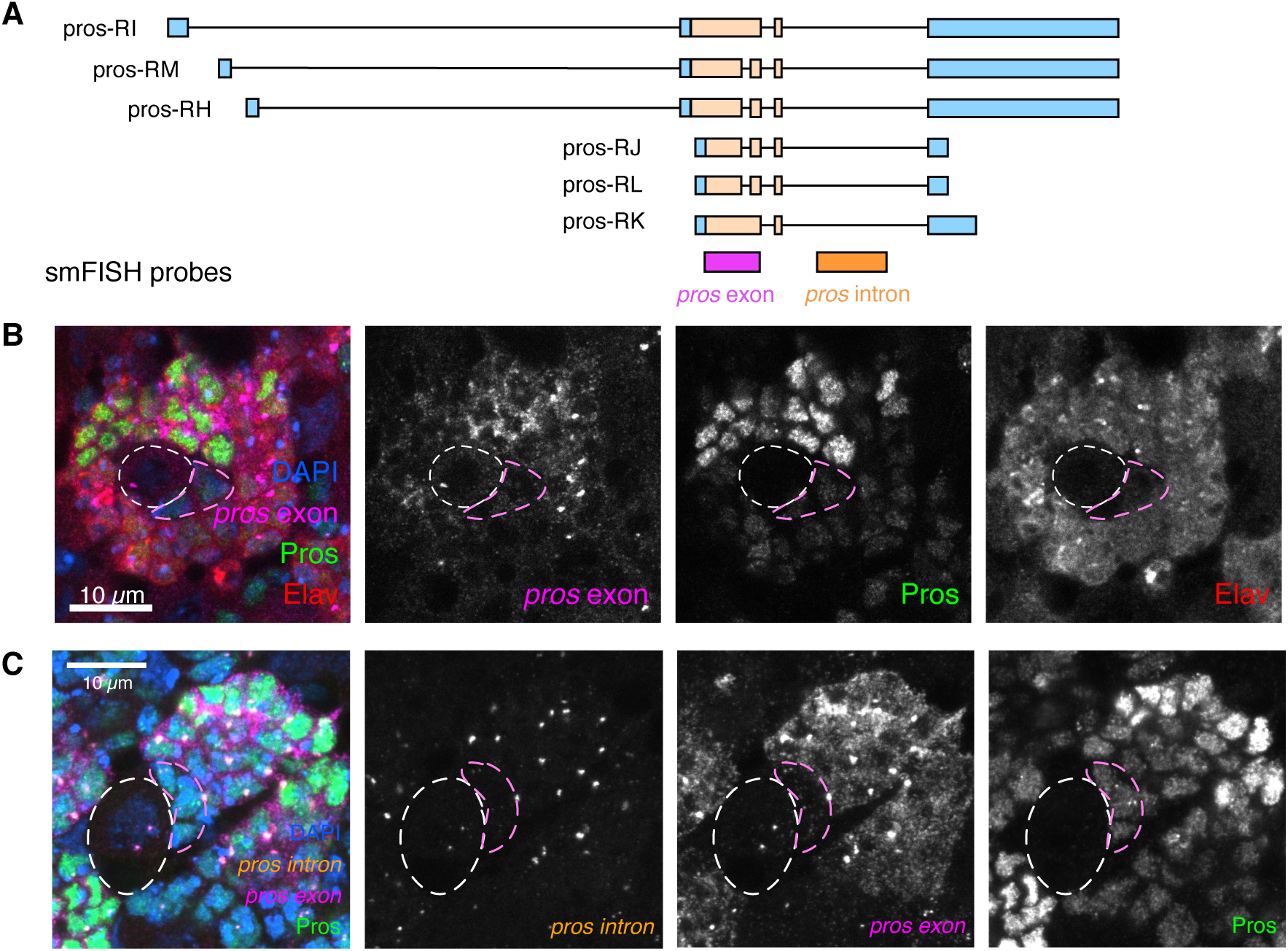
Pros cell type-specific expression is regulated post-transcriptionally. **(A)** Illustration of smFISH probe positions on the *pros* mRNA transcript isoforms. **(B)** Third instar larval brains stained for smFISH against *pros* exon, with IF against Elav and Pros. *pros* mRNA is upregulated in post-mitotic neurons (marked with Elav), and this correlates with upregulated Pros protein. (**C**) smFISH against the *pros* intron and *pros* exon (probe positions shown in **A**) together with IF against Pros protein, shows that *pros* is transcribed in all cells of the Type I NB lineage, even in NBs where there are low levels of Pros protein and *pros* mRNA. Scale bar: 10 µm. NB: White dashed outline, GMC: pink outline

We then asked whether the increase in *pros* mRNA levels in neurons is regulated at the level of transcription, or post-transcriptionally through mRNA stability. We used smFISH with *pros* intron-specific probes to visualise the *pros* transcription foci (Figure 1A, C). There are two transcription foci in each cell (these are paired in the progeny cells, thus visible as a single combined focus of higher intensity). Unexpectedly, we observed bright transcription foci in the type I NBs, but very few cytoplasmic transcripts in these cells (Figure 1C, white outline). This result suggests that *pros* mRNA is more unstable in NBs compared to neurons, where much higher levels of *pros* mRNA accumulate.

### *pros* mRNA in larval brains contains an exceptionally long 3’ UTR

Post-transcriptional regulation is often linked to expression of distinct alternative isoforms, and neurons have been shown to selectively express transcript isoforms with unusually long 3’ UTRs (Hilgers et al., 2012; Hilgers et al., 2011; Oktaba et al., 2015). Therefore, we examined the *pros* isoform usage in the type I NB lineage.

*pros* is annotated in FlyBase as having six isoforms with three different 3’ UTR lengths: ∼1 kb, ∼3 kb and ∼15 kb (Figure 2A, FB2019_05). To test which isoforms are expressed in larval brains, we carried out Northern blots. Our Northern blot analysis using a probe that is common to all the annotated isoforms of *pros* transcripts (probe ALL) detected the known ∼6 kb isoform in embryos (Figure 2A, B) (Doe et al., 1991; Matsuzaki et al., 1992; Vaessin et al., 1991). In larval brain extracts, the Northern blots show multiple-sized *pros* transcripts including isoforms approximately 6 kb, 9 kb and 21 kb in length (Figure 2B), corresponding to 3’ UTR lengths of 1 kb, 3 kb and 15 kb, respectively. We also detected the 21 kb *pros* band in larval brains using a probe targeting the extended region of the 15 kb 3’ UTR (Figure 2A, B). We conclude that embryos express only the shortest isoform of *pros*, whereas larval brains express two additional isoforms, with 3 kb and 15 kb long 3’ UTRs, respectively.

**Figure 2:**
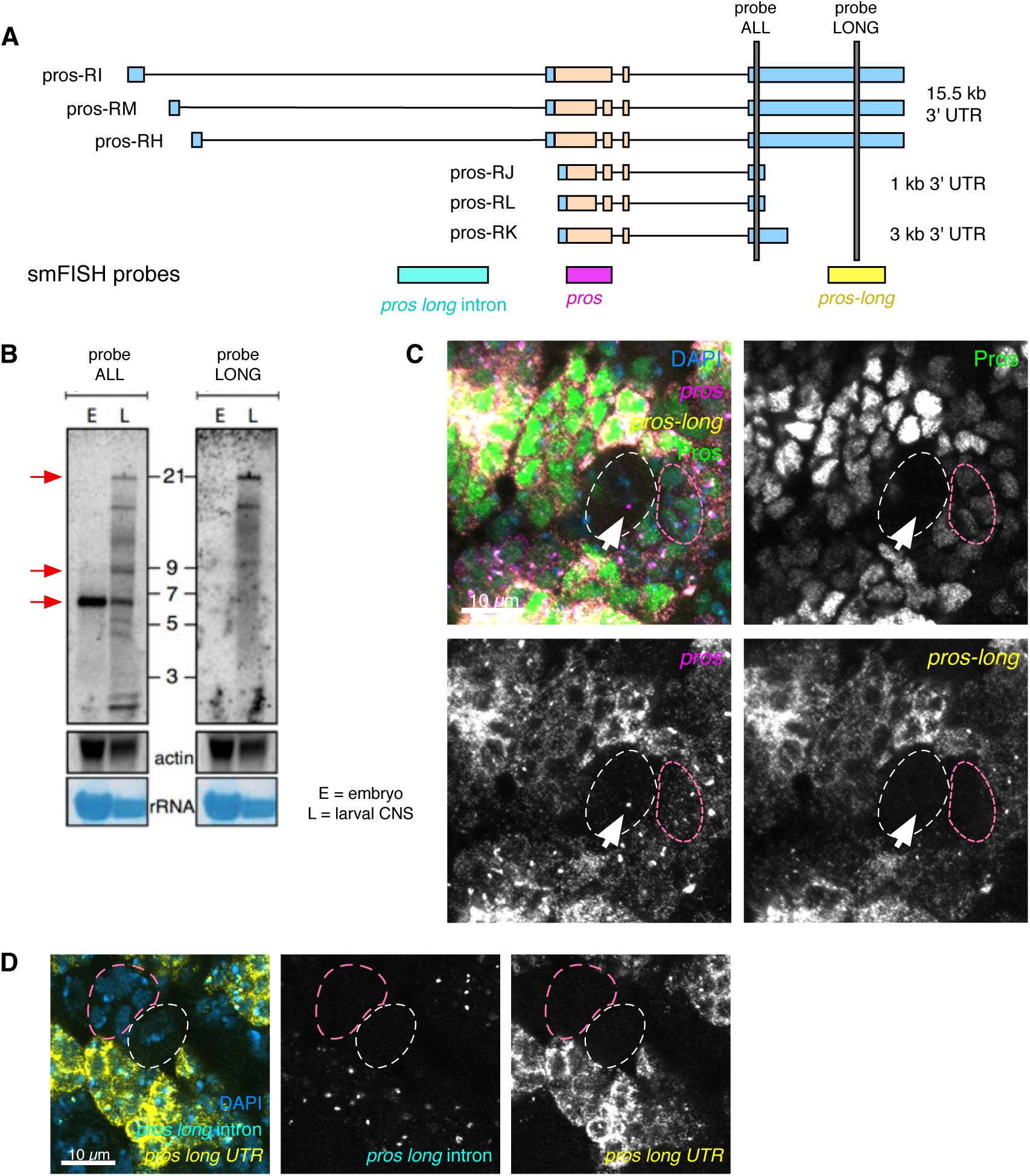
*pros^long^* isoform is expressed brains and is specifically transcribed in larval neurons. **(A)** Diagram showing annotated *pros* transcript isoforms and the position of smFISH (*pros long* intron, *pros* and *pros-long*) and Northern blot probes (grey bars). **(B)** Northern blot confirms the existence of multiple *pros* mRNA isoforms at 6 kb, 9 kb and 21 kb (red arrows) in the larval CNS. The 21 kb isoform corresponds to *pros* transcript with 15 kb 3’ UTR. E: embryo, L: larval CNS **(C)** Costaining with *pros* exon and *pros-long* smFISH and Pros IF shows that *pros^long^* RNA is detected only in neurons. In the NB (white dotted outline), or the GMCs (pink dotted outline), *pros* exon is detected without *pros-long*. Pros protein is upregulated in cells expressing *pros^long^*. **(D)** *pros long* intron probe only recognises the nascent transcripts of the *pros^long^* isoform. *pros long* intron is detected exclusively in neurons, and not in the NB (white dotted outline) or GMCs (pink dotted outline). Scale bar: 10 µm.

To investigate the cell type-specific expression of the *pros* isoforms in larval brains, we performed smFISH experiments using a probe specific to the shared coding exon (*pros)*, and a probe that is specific to the extended region of the 15 kb 3’ UTR region (*pros-long*) of the *pros^long^* transcripts (defined as the isoforms containing the 15 kb 3’ UTR), co-stained for Pros protein (Figure 2A,C). *pros^long^* is highly expressed in the cells with highest *pros* exon signal, and correlates well with upregulated Pros protein. We conclude that the upregulated *pros* mRNA in neurons consists primarily of *pros^long^.* Co-staining with Elav to label neurons shows that *pros^long^* is expressed specifically in neurons and is absent from NBs and GMCs (Figure S1A, GMCs dashed outline).

However, GMCs do show low levels of *pros* exon smFISH signal and Pros protein (Figure 2C, outlined in dashed pink). These observations suggest that the short isoform of *pros* produces the low levels of Pros seen in the GMCs and NBs.

To determine precisely where *pros^long^* is transcribed in the brain, we visualised *pros^long^* transcription using smFISH with intron probes specific for the long isoforms (Figure 2A, D). We detected *pros^long^*-specific transcription foci only in neurons, where we also observed *pros-long* smFISH signal in the cytoplasm. In contrast, no *pros-long* intron signal was detected in the NBs or GMCs (Figure 2D outlined in dashed white and pink respectively). We conclude that the *pros^long^* isoform is specifically transcribed in larval neurons. Our results support a model in which a low level of Pros protein is produced by the short *pros* isoform in NBs and GMCs. Transcription of *pros^long^* is switched on in the larval neurons and the isoform is stabilised, resulting in higher *pros* mRNA levels and upregulated Pros protein.

### Syp is expressed in the type I NB lineage and stabilises *pros* mRNA

Post-transcriptional regulation, including mRNA stability, depends on the binding and action of specific RBPs. We have previously shown in larval extracts that *pros* mRNA associates with Syp, a highly conserved RBP (McDermott et al., 2014). Syp is a key factor in the temporal regulation of larval NBs and is known to regulate the transcription factor Chinmo to determine neuron fate (Liu et al., 2015; Ren et al., 2017; Syed et al., 2017). Syp is expressed in the larval type I NBs and their progeny (Figure 3A) (Liu et al., 2015), and so we tested whether Syp binds *pros* mRNA in the larval brain. 58.3% of *pros* mRNA was co-immunoprecipitated with Syp in larval brain lysates, while only 3.2% of *rp49,* a control RNA, was pulled down (Figure 3B, C). These results demonstrate that Syp binds specifically to *pros* RNA, rendering Syp a good candidate for an upstream regulator of *pros* mRNA stability.

**Figure 3:**
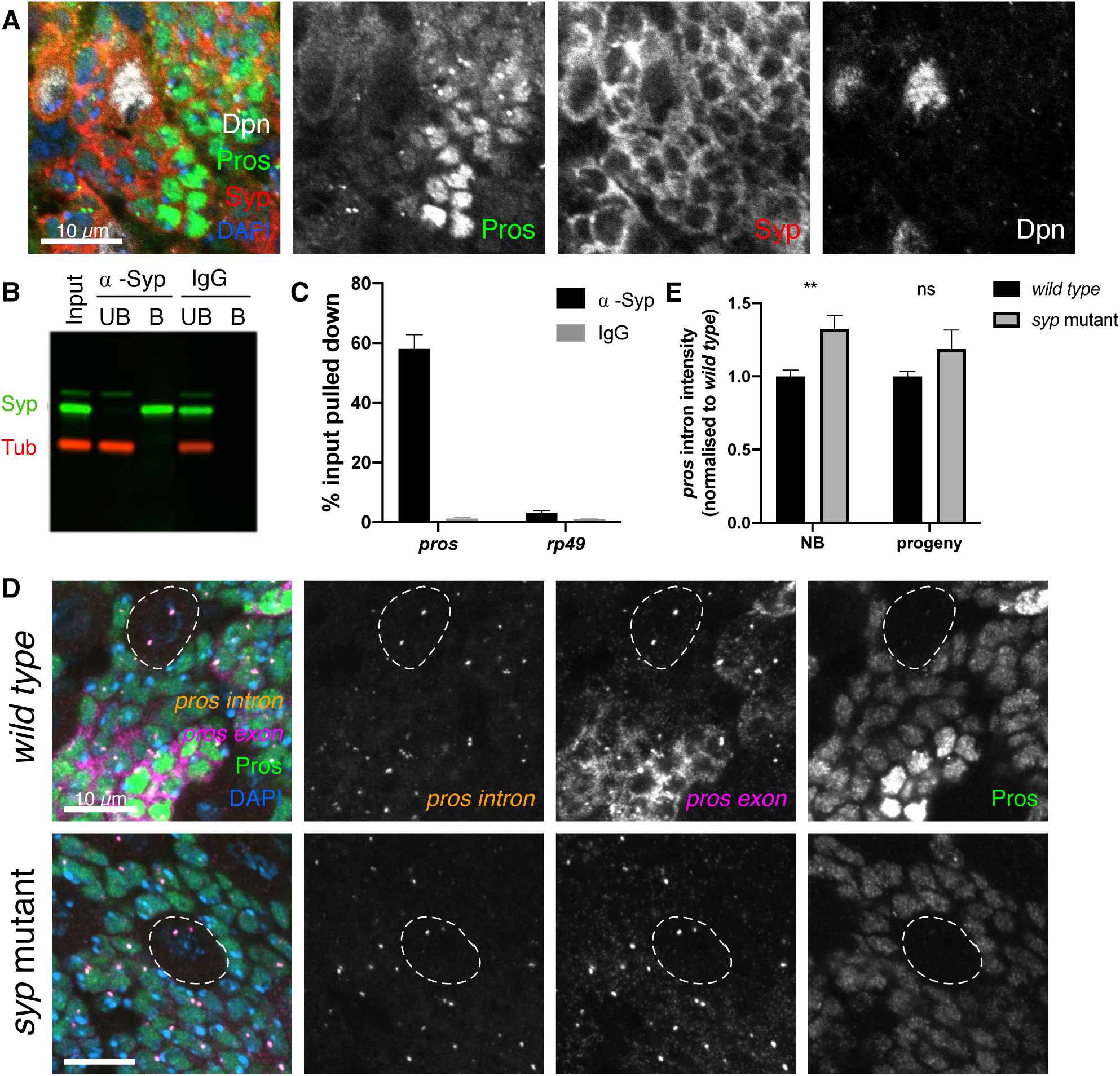
Syncrip selectively stabilises the *pros^long^* isoforms in neurons. **(A)** Syncrip (Syp) protein is expressed in the type I NB (marked with Dpn) and progeny lineage in 3^rd^ instar larvae. Syp is also expressed in the neurons, where Pros is upregulated. **(B)** Western blot confirms Syp is immunoprecipitated (IP) selectively and efficiently. ⍺-Tubulin (Tub) was used as negative control. **(C)** *pros* mRNA is enriched by Syp IP (58.3% of input pulled down) but not by IgG control IP (1.3%). Negative control, ribosomal *rp49 (*⍺-Syp: 3.2%, IgG: 0.9%) (*n* = 7). Error bars represent SEM. **(D)** Loss of Syp has a minimal effect on transcription levels indicated by *pros* intron, but *pros* exon signal and Pros protein is significantly reduced. NBs outlined in white. **(E)** Intensity of *pros* intron signal in *wild type* and *syp* mutant. Normalised to *wild type* for each experiment. NB: sum of intensity of two transcription foci, progeny: intensity of single spot including both transcription foci. Scale bars: 10 µm

To determine whether Syp regulates the levels of *pros* mRNA, we compared levels of *pros* transcription, cytoplasmic *pros* transcripts and Pros protein in brains mutant for a *syp* allele that lacks Syp protein (Methods). The results show that Pros protein is reduced in the neurons of *syp* mutant brains, to similar levels to those found in *wild type* GMCs (Figure 3D). Pros protein levels are maintained at low levels in the *syp* mutant neurons but the upregulation of Pros is lost. *pros* mRNA is also very substantially reduced in the *syp* mutants, most obviously in the neurons (Figure 3D). We measured the intensity of *pros* intron signal at the transcription foci in *wild type* and *syp* mutant brains (Figure 3E) and observe a small but significant increase in *pros* transcription in the *syp* mutant type I NBs, compared to *wild type* (1.3-fold increase). In progeny cells (pooled neurons and GMCs) there is no significant change in *pros* transcription in the *syp* mutant, compared to the *wild type* (Figure 3E). Therefore changes in *pros* transcription are not responsible for the reduction of *pros* transcripts in the *syp* mutant. We suggest that Syp mediates the upregulation of Pros protein in neurons primarily by stabilising cytoplasmic *pros* mRNA.

Syp negatively regulates the RNA-binding protein Imp/IGF2BP in type I NBs (Liu et al., 2015) and therefore could regulate *pros* mRNA stability indirectly via Imp. To distinguish whether Syp regulates *pros* directly or indirectly, we compared the levels of *pros* mRNA and Pros protein in *syp* knockdown and *imp syp* double-knockdown brains (Yang et al., 2017a) (Figure S1B). *imp syp* double-knockdown phenocopies the single *syp* knockdown (Figure S1B). Combined with the *pros* pulldown in Syp immunoprecipitations, these results indicate that Syp’s effects on *pros* are independent of its effects on Imp.

### *pros^long^* is stabilised by Syp

Loss of Syp has the most significant effect on *pros* in the neurons. In *syp* mutant neurons *pros* mRNA is substantially reduced and Pros protein upregulation is lost. We previously showed that *pros^long^* is responsible for the neuronal *pros* mRNA and protein upregulation, so we asked whether Syp acts specifically on *pros^long^.* We used smFISH against the *pros^long^* 3’ UTR in *syp* mutant larval brains and found that it is significantly reduced in neurons, compared to *wild type* (Figure 4A). We co-stained with the *pros^long^*-specific intron probe and found that *pros^long^* is transcribed normally in the *syp* mutant. We conclude that Syp upregulates Pros protein in neurons by stabilising *pros^long^* mRNA. To confirm that the loss of cytoplasmic *pros* mRNA in *syp* mutants is due to specific loss of the long isoform, we carried out transcriptomics analysis of *syp* mutant versus *wild type* brains. The results show that the abundance of *pros^long^* is specifically reduced in *syp* mutant brains, whereas the shorter *pros* forms are maintained (Figure 4B, C). Northern blot analysis confirmed these findings (Figure 4D).

**Figure 4:**
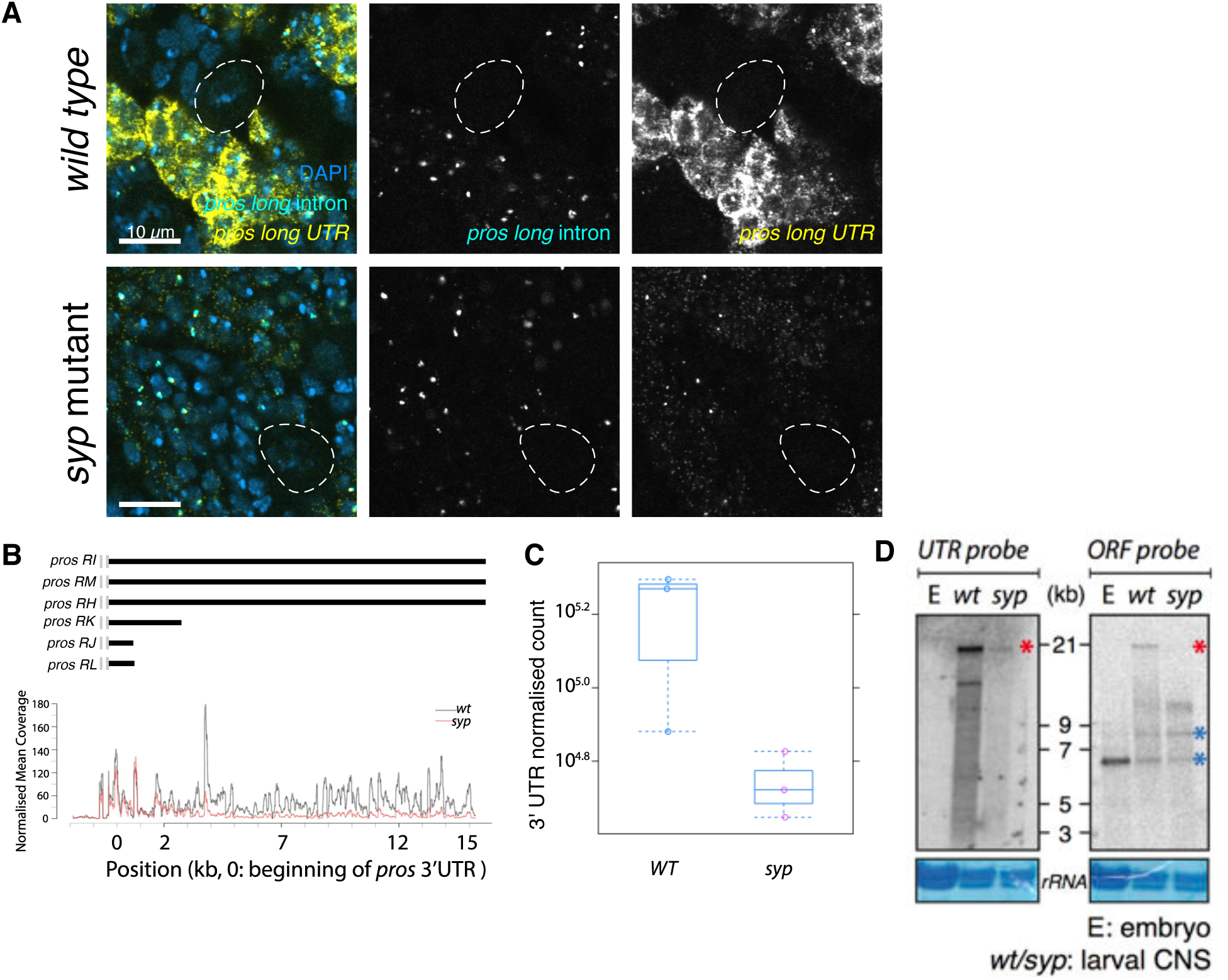
Syncrip selectively stabilises the *pros^long^* isoforms in neurons. **(A)** smFISH against *pros long* intron and *pros long* shows that *pros^long^* is normally transcribed in the *syp* mutant. **(B)** RNA sequencing shows selective loss of *pros^long^* isoform in *syp* mutants. Read coverage for the first 3 kb of *pros* 3’ UTR is not different between *wild type* (wt) and *syp* mutant whereas the distal 12 kb of *pros* 3’ UTR is reduced in *syp* mutants. **(C)** Quantitative analysis of total sequencing reads of *pros* 3’ UTR in *wild type* and *syp* mutants. **(D)** Northern blot showing the selective loss of *pros^long^* isoforms in *syp* mutants. The intensity of the 6.5 kb and 9 kb bands are similar between *wild type* and *syp* mutants (blue asterisks) whereas the 21 kb band is significantly reduced in *syp* mutants (red asterisk).

### *pros^short^* is sufficient to maintain neuronal fate

We have shown that Syp is required for the upregulation of Pros protein in larval neurons, and this occurs through the regulated mRNA stability of *pros^long^*. When Pros is depleted from neurons, but not NBs or GMCs, such as in the *midlife crisis* mutant, this leads to loss of neuronal differentiation and expression of NB genes (Carney et al., 2013). Therefore, we used the *syp* mutant as a tool to examine the specific role of the upregulated Pros protein levels in neurons, as opposed to the effect of loss of all Pros protein from the neurons. Unexpectedly, we found that neuronal differentiation progresses normally in *syp* mutants and neurons do not revert to NBs, as they do not switch on the NB marker, Dpn. We used clonal analysis to mark individual NB lineages in *wild type* and *syp* mutant brains and stained with Dpn, Pros and Elav (Figure 5). Each *syp* mutant clone contained just one Dpn +ve NB, and Elav was expressed in the progeny cells, demonstrating their neuronal identity. We conclude that the stabilisation of *pros^long^* by Syp in progeny cells is not required for the normal neuronal path of differentiation nor to prevent reversion of neurons to NBs. We conclude that the remaining low level of Pros in the neurons of the *syp* mutant, provided by the short *pros* mRNA isoform, is sufficient to prevent reversion to a NB fate.

**Figure 5:**
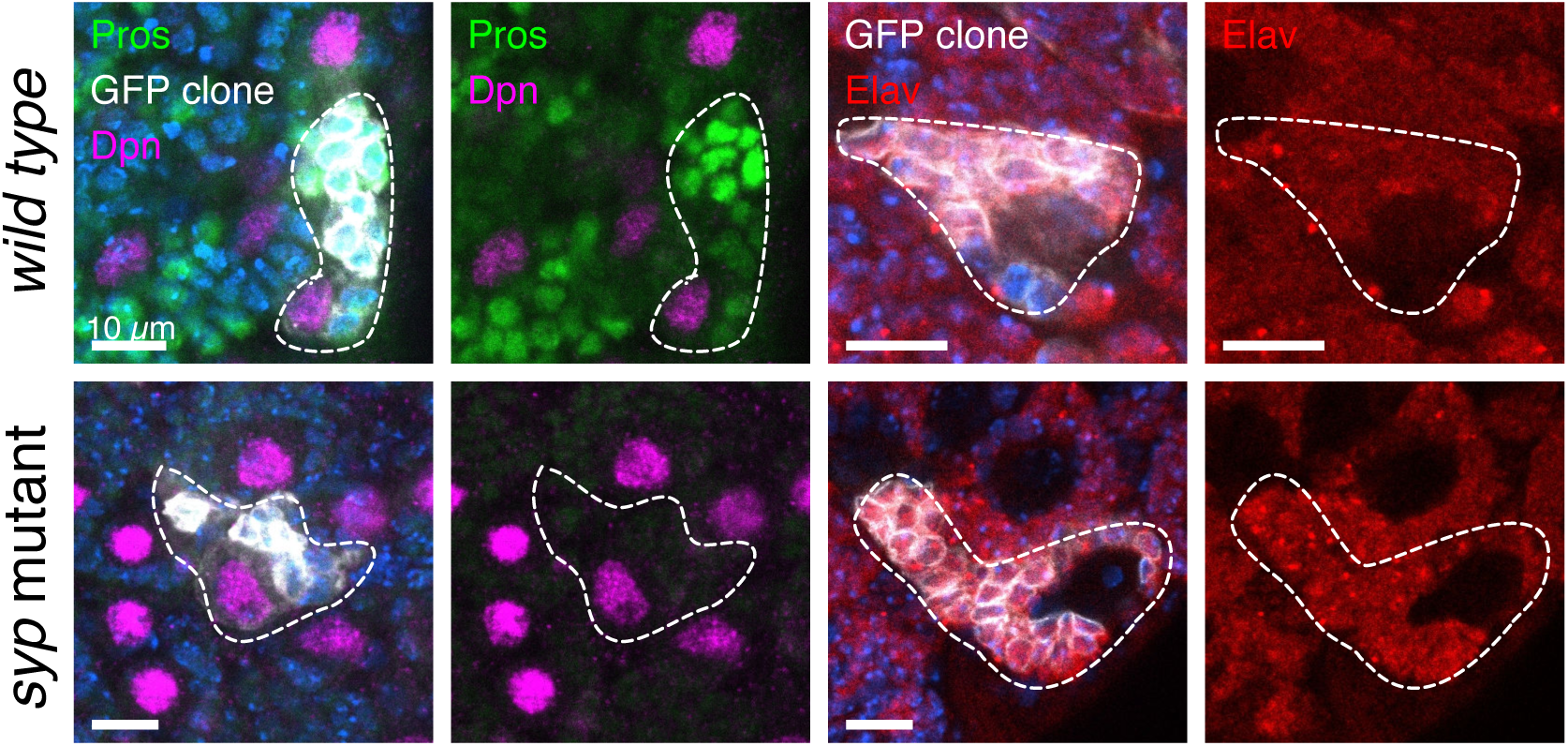
Syp and *pros^long^* are not required to maintain larval neurons in differentiated states. Clonal analysis shows normal GMC differentiation in the *syp* mutant. Type I NB lineage clones are marked with mCD8::GFP (outlined with white trace) and stained with IF against Pros, Dpn (wt n = 7, *syp* n = 9) and Elav (wt n = 16, *syp* n = 4). Both *wild type* and *syp* mutant clones have a single Dpn-expressing cell per clone and the progeny cells express Elav, marking a neuronal fate.

The question remains: what is the function of *pros^long^* mRNA? A burst of nuclear Pros protein is known to have an important role in terminating NB divisions in pupae (Kohwi and Doe, 2013; Maurange et al., 2008). Furthermore Syp is required for the final symmetric division that terminates the type I NB in the pupa (Yang et al., 2017a). In the wild type pupal brain, all type I NBs, except the mushroom body NBs, terminate by 48 h after pupal formation (APF) (Siegrist et al., 2010). Syp-depleted type I NBs persist in the pupal brain for more than 48 h APF (Yang et al., 2017a). We surmised that *pros^long^* might be required at the terminal division, stabilised by Syp to allow the upregulation of Pros in the NB. To test this hypothesis we produced a series of transgenic lines aiming to remove *pros^long^*.

Several attempts to generate flies selectively deleting the *pros^long^* transcripts were unsuccessful. These included a frame shift mutation, which did not introduce nonsense-mediated mRNA decay (NMD) of the long *pros* isoforms (Figure S2, Methods), and attempting a direct deletion of the *pros* 3’ UTR extension (Methods). We successfully deleted the three promoters that are annotated to produce *pros^long^* (Figure S3A), but found that the 3’ UTR extension of *pros^long^* can be transcribed from the downstream promoters (Figure S3B) and there was no change in Pros protein expression (Figure S3C). Finally, we used a strong SV40 transcriptional terminator to terminate transcription at the beginning of the 3’ UTR extension (Methods). The SV40 terminator reduced but did not entirely remove the extended UTR of *pros^long^* (Figure S4). Unexpectedly, the inserted *dsRED* marker used to screen for integration does terminate the 3’ UTR extension of *pros* (Figure 6), allowing us to test the effect of removing the 3’ UTR extension. The *dsRED* inserted in the opposite orientation to *pros* (*proslong-REDr*) gave the lowest background and was used in further experiments (Methods).

**Figure 6:**
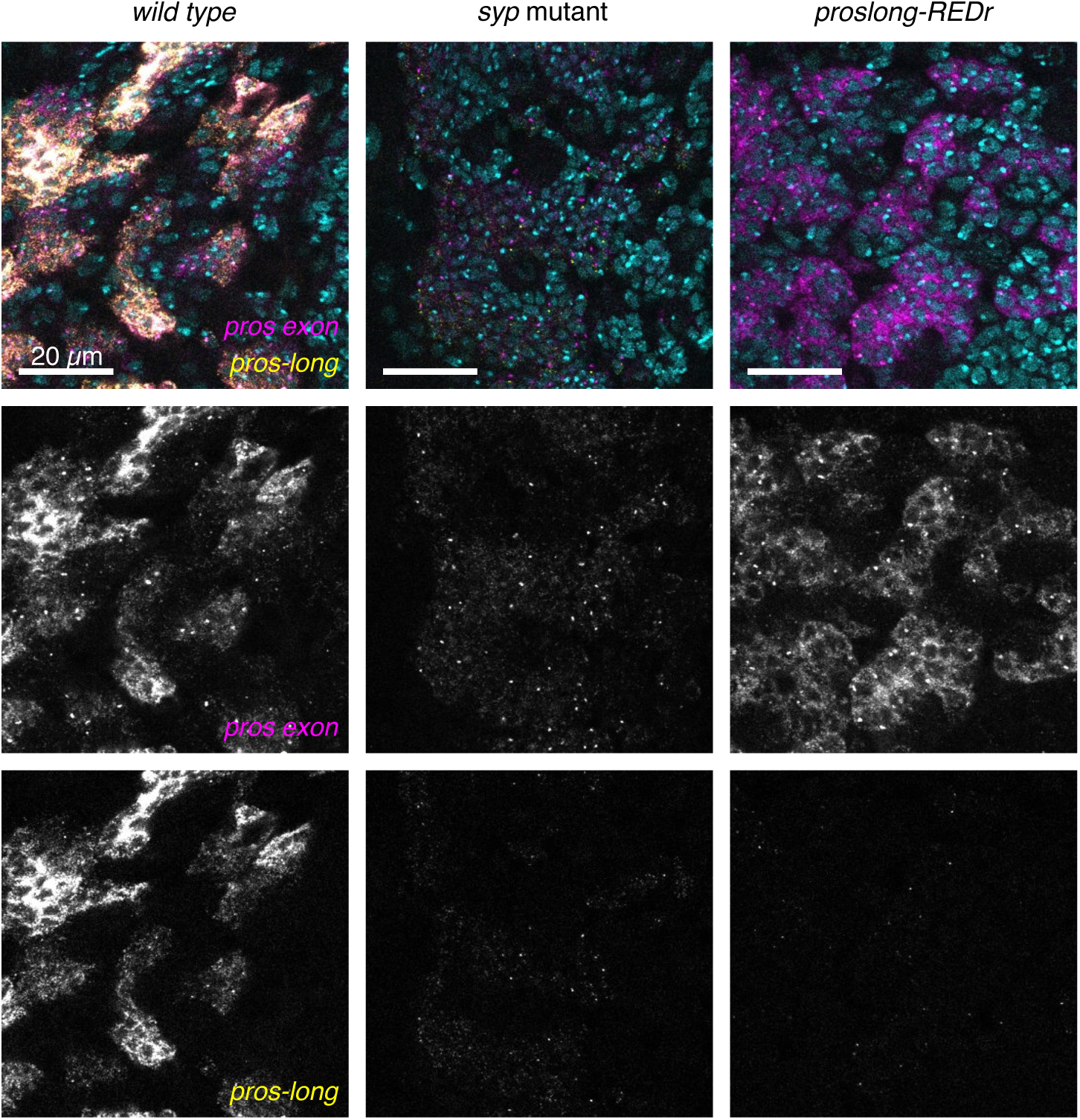
*proslong-REDr* terminates transcription of the *pros* UTR extension. Insertion of a dsRED marker in reverse orientation downstream of the 3 kb UTR termination signal (*proslong-REDr*) results in loss of the *pros* UTR extension (visualised with the *pros-long* smFISH probe). The *pros exon* smFISH shows that *proslong-REDr* expresses higher levels of *pros* than *syp* mutant brains, but much less than *wild type*.

smFISH against the *pros* exon and *pros-long 3’ UTR* showed that the long 3’ UTR is almost entirely lost in the *proslong-REDr* brains (Figure 6). Therefore, regulation of *pros* mRNA through the 3’ UTR extension is disrupted in this line. The signal from the *pros* exon probes is reduced compared to the *wild type,* although significantly higher than in the *syp* mutant, which could be explained by residual binding of Syp to *pros^long^* in regions outside the UTR extension. Alternatively, Syp may stabilise multiple isoforms of *pros*.

In order to understand the role of *pros^long^* in regulating Pros protein expression, we examined Pros protein level in *proslong-REDr* brains. We found that the cell type-specific upregulation of Pros protein in neurons is lost in *proslong-REDr* brains (Figure 7A), consistent with the 3’ UTR extension of *pros* being required for the neuronal upregulation of Pros protein during larval neurogenesis.

**Figure 7:**
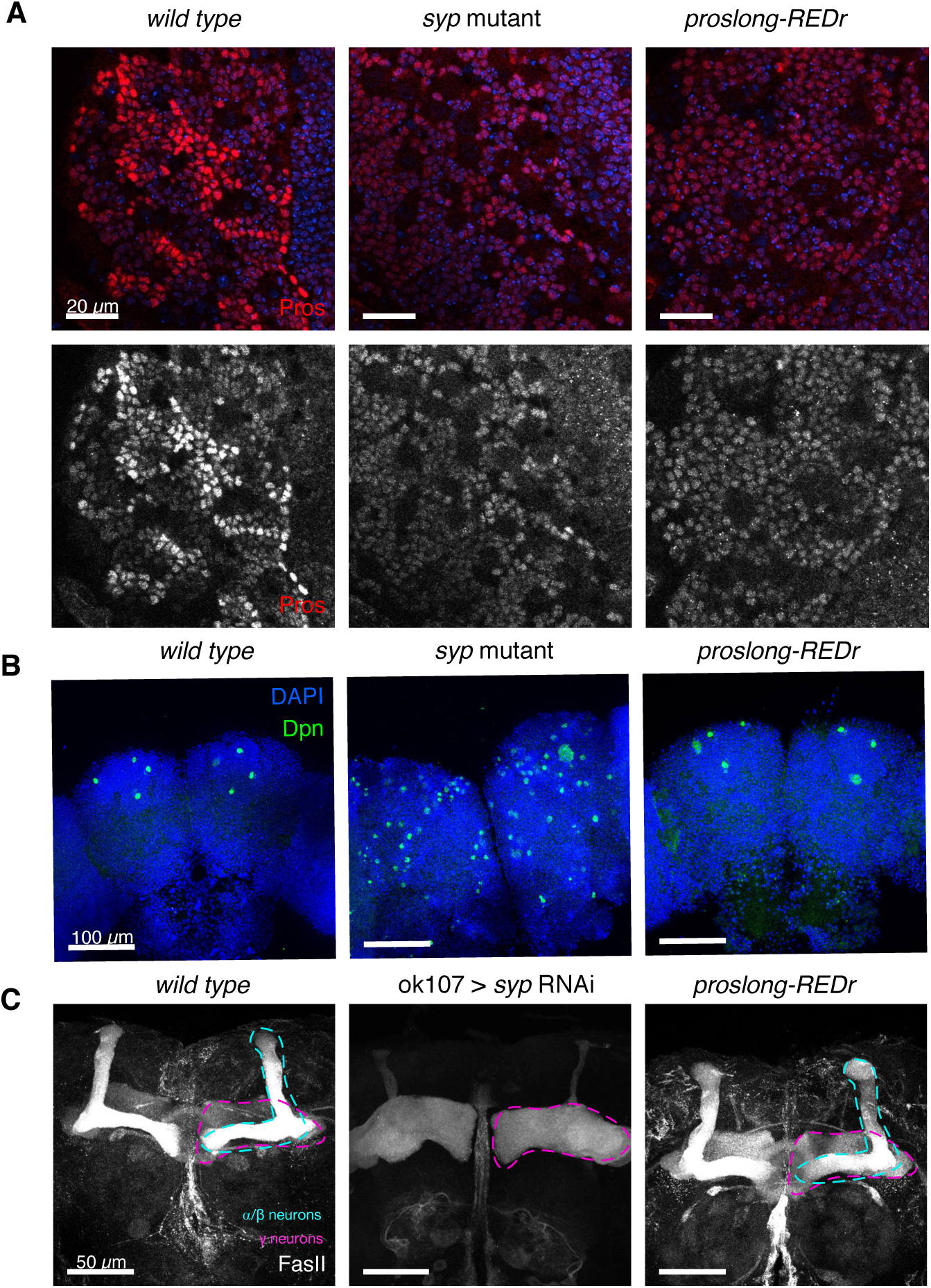
Loss of *pros^long^* blocks Pros protein upregulation but has no effect on NB termination A. Pros protein level is not upregulated in the neurons of *proslong-REDr* larval brains. **B** Dpn marks NBs in pupal brains, 48 hr APF. In *wild type*, only the four MB NBs remain. In the *syp* mutant, many NBs are long-lived in the pupal brain. In the *proslong-REDr* brains, type I NBs terminate normally and only the four MB NBs persist. **C** FasII IF stains γ and α/β neurons in the mushroom body (MB) of 1 day old adult brains. There is no difference in MB morphology in the *proslong-REDr* line, compared to *wild type*.

To test the function of *pros^long^* in NB termination, we examined the number of NBs remaining at 48 hr APF. We found that *pros^long^* is not required for normal NB termination. At 48 hr APF, *wild type* pupae have only four NBs remaining (labelled with Dpn, the mushroom body (MB) NBs), while *syp* mutant pupal brains contain many persistent NBs (Figure 7B). In the *proslong-REDr* brains, we observe only the four MB NBs at 48 hr APF, showing that the type I NBs terminate normally, despite the loss of the *pros^long^* 3’ UTR extension.

We tested whether *pros^long^* is required for neuron fate specification. Syp controls α/β neuronal fate during mushroom body (MB) development, and loss of Syp results in complete loss of α/β neurons and precocious production of γ neurons (Liu et al., 2015). Single-cell transcriptomics analysis of adult *Drosophila* brain reveals that α/β neurons express much higher levels of *pros* compared to γ and ⍺’/β’ neurons (Davie et al., 2018). To test whether stabilisation of *pros^long^* by Syp is required for the specification of α/β neurons, we examined the MB morphology in 1-day old *proslong-REDr* adult brains (Figure 7C). Immunostaining against FasII showed the overall morphology of α/β and γ neurons in the *proslong-REDr* brains are comparable to the *wild type* control, indicating that fate patterning of MB neurons occurs normally even in the absence of *pros^long^*.

Finally, we wondered whether *pros^long^* we also tested whether *pros^long^* is required in adult flies. We found that adult flies lacking *pros^long^* show highly penetrant behavioural defects. The majority of *proslong-REDr* adult flies eclose normally and are morphologically normal, but display an abnormal behaviour consistent with impaired locomotor or neurological activity (Movie S1). This phenotype indicates that the *pros^long^* mRNA isoform has a function in the adult brain, although its detailed characterisation is beyond the scope of this study.

## DISCUSSION

Many key regulators of NB proliferation and differentiation have now been identified and characterised (Grosskortenhaus et al., 2006; Isshiki et al., 2001; Kambadur et al., 1998; Kohwi and Doe, 2013; Li et al., 2013; Narbonne-Reveau et al., 2016; Novotny et al., 2002; Pearson and Doe, 2003). Recently an increasing number of RBPs, the key regulators of post-transcriptional processes, have been implicated in neurodevelopment (Betschinger et al., 2006; Hilgers et al., 2012; Liu et al., 2015), suggesting that the importance of post-transcriptional regulation in the brain has so far been underestimated. Here, we have applied smFISH to examine the regulation of the transcription factor (TF) Pros, a master switch promoting neuronal differentiation. We show that Pros expression is regulated by the differential stability of its mRNA isoforms depending on their 3’ UTR lengths. Unstable *short* mRNA isoforms produce sufficient Pros protein to prevent GMCs and their neuronal progeny from reverting back to NB identity, but a switch to the more stable *pros^long^* isoforms is required to upregulate Pros protein in larval neurons. Our findings highlight the capacity of cell type-specific alternative 3’ UTRs to mediate different modes of post-transcriptional regulation of mRNA isoforms.

### Low level *pros^short^* has a distinct function from high level *pros^long^*

Surprisingly, we found that neurons do not de-differentiate into NBs in *syp* mutants, despite the loss of neuronal Pros upregulation in the absence of *pros^long^*. Previous work has shown that Pros elimination in young or middle-aged larval neurons causes de-differentiation of neurons and their reversion to NBs (Bello et al., 2006; Betschinger et al., 2006; Carney et al., 2013; Lee et al., 2006). Our work suggests that the low levels of Pros remaining in the *syp* mutant (provided by *pros^short^*) are sufficient for neurons to maintain their identity. *pros^long^* is not required to drive differentiation of GMCs to neurons or to maintain neuronal identity. Although we have not uncovered the function of *pros^long^* in larval/pupal brain development, the impaired locomotive activity of the *proslong-REDr* adult flies indicates a role of *pros^long^* in the adult brain, either because of an earlier neuronal specification event or due to a function of Pros in the adult brain.

### The 15 kb *pros* UTR extension allows multiple levels of post-transcriptional regulation

Exclusion of the 3’ UTR extension from *pros* transcripts in the *proslong-REDr* brains, reduces the number of *pros* transcripts and Pros protein in the neurons. However, the *pros* transcript levels are still much higher in *proslong-REDr* than *syp* mutant brains. This residual *pros* signal suggests that Syp can stabilise *pros* mRNA through binding to additional regions of the transcript, perhaps the 5’ UTR sequence that is also unique to the *pros^long^* transcripts, or some shared sequence included in all transcripts.

The 3’ UTR extension of *pros^long^* may mediate a second regulatory step, at the level of translation. The upregulation of Pros protein in the neurons is lost in the *proslong-REDr* brains, despite the relatively high levels of remaining *pros* mRNA transcripts. While the *pros* exon smFISH signal is much higher in the *proslong-REDr* brains, compared to the *syp* mutant, the Pros protein levels are similar between the two genotypes. This result suggests that the *pros^long^* 3’ UTR extension includes additional regulatory sequences that promote increased *pros* translation, either via Syp or an unidentified second RBP. This hypothesis would explain why a moderate decrease in *pros* transcript levels in *proslong-REDr* brains leads to a complete loss of neural Pros protein upregulation.

### Post-transcriptional and transcriptional regulation work hand-in-hand to achieve cell type-specific gene expression

Our experiments show that Pros expression is controlled at two levels: alternative polyadenylation and then differential mRNA stability, regulated through Syp binding. The molecular mechanism underlying the cell type-specific choice of *pros* isoform has not yet been identified. In *Drosophila* embryos, Elav is recruited at the promoter of extended genes and is required to extend the 3’ UTR of *brat* (Hilgers, 2015; Oktaba et al., 2015). Future experiments will determine whether *pros* differential polyadenylation is regulated at the promotor region by a similar Elav-dependent mechanism.

Many key regulators in the brain also have complex gene structures such as multiple isoforms and long 3’ UTRs, hallmarks of post-transcriptional mechanisms (Berger et al., 2012; Hilgers et al., 2011; Stoiber et al., 2015; Tekotte et al., 2002). Such genes include the temporal regulator neuronal fate, *chinmo* (Liu et al., 2015), driver of cell growth and division, *myc* (Samuels et al., 2019) and the mRNA-binding proteins, *brat* (Bello et al., 2006; Betschinger et al., 2006; Lee et al., 2006) and *imp* (Liu et al., 2015). Quantitative smFISH approaches combined with genetics and biochemistry will allow the detailed disentanglement of transcriptional and post-transcriptional mechanisms regulating these genes.

### Regulating expression levels by differential mRNA isoform stability and long 3’ UTRs is likely to be a conserved mechanism

Mammalian SYNCRIP/hnRNPQ, is an important regulator of neural development (Lelieveld et al., 2016; Liu et al., 2015; Stoiber et al., 2015), and has a number of post-transcriptional roles including regulating mRNA stability through binding at the 3’ end of transcripts (Kim et al., 2011; Kuchler et al., 2014). Prox1, the mammalian orthologue of Pros, is a tumour suppressor that regulates stem cell differentiation in the brain as well as many other organ systems (Elsir et al., 2012; Stergiopoulos et al., 2014). Like *Drosophila pros*, *prox1* has several isoforms including alternative 3’ UTRs, and a burst of Prox1 expression is required to drive the differentiation of immature granular neurons in the adult hippocampus (Hsieh, 2012). It is also likely that post-transcriptional regulation by RBPs helps determine the expression and translation of Prox1. Application of quantitative smFISH to mammalian systems will uncover whether Prox1 expression levels, like *pros,* are regulated through differential stabilisation of different 3’ UTR isoforms.

## AUTHOR CONTRIBUTIONS

Conceptualisation, T.J.S., L.Y., D.I.H. and I.D.; Investigation, T.J.S, Y.A. (Northern blots), J.Y.L. (adult mushroom body morphology and adult phenotyping), L.Y. (RNA preparation for Northern blots, making the NMD deletion); Genomics analysis, A.I.J.; Data Analysis, T.J.S., Y.A., A.I.J., J.Y.L.; Experimental design, T.J.S, Y.A., J.Y.L., F.R. (help and advice on all CRISPR constructs), L.Y., C.P.Y., T.L. D.I.H., I.D.; Writing -Original Draft, T.J.S., L.Y.; Writing - Review & Editing, T.J.S., D.I.H., I.D.; Supervision, I.D.

## ACKNOWLEDGEMENTS

We are grateful to Andrew Bassett (Genome Engineering Oxford, GEO) for assistance in designing and validating CrispR guide RNAs, Tomek Dobrzycki for initial work that lead to the project, MICRON Oxford (http://micronoxford.com, supported by a Wellcome Strategic Awards to ID (091911/B/10/Z and 107457/Z/15/Z) providing access to equipment and advice on advanced imaging techniques, Alan Wainman and Richard M. Parton for advice on advanced microscopy, Darragh Ennis for help and advice with fly husbandry and administration. We are also indebted to Alfredo Castello, Lidia Vasilieva, Jordan Raff and Neil Brockdorff for their comments. We would also like to thank the Cambridge Fly Facility for transgenic production and the Bloomington *Drosophila* Stock Centre. T.J.S. was funded by Wellcome Trust Four-Year PhD Studentship (105363/Z/14/Z) and Wellcome Investigator Award 209412/Z/17/Z. L.Y. was funded from the Clarendon Trust and Goodger Fund. F.R. was funded by a Marie Curie Postdoctoral Fellowship and Wellcome Investigator Award 209412/Z/17/Z. C.P.Y. and T.L. were supported by Howard Hughes Medical Institute. J.Y.L. was funded from the Clarendon Trust. D.I.H. was funded by University College London. I.D. and A.I.J were funded from a Wellcome Senior Research Fellowship (096144/Z/17/Z) and Wellcome Investigator Award 209412/Z/17/Z.

## SUPPLEMENTARY INFORMATION

**Figure S1:**
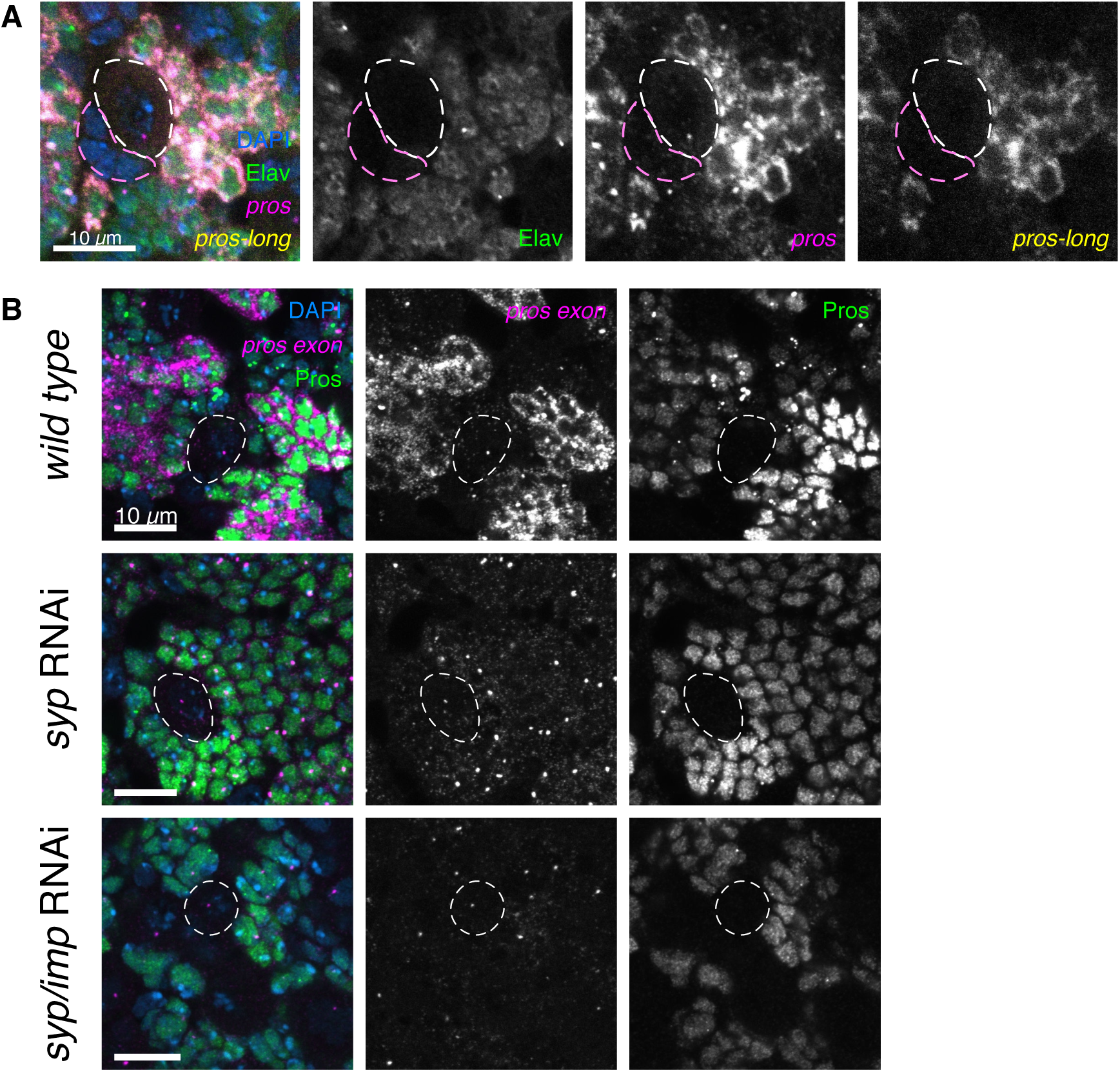
Syp stabilises *pros* directly, not via Imp. **A** Staining with Elav IF (marking post-mitotic neurons), and *pros* exon and *pros-long* smFISH. *pros^long^* is only expressed in Elav+ cells, and is not expressed in the NBs (white outline) or GMCs (pink outline) **B** Brains stained with Pros protein and *pros* exon smFISH. *pros* is lost in the *syp* RNAi knockdown, and in the *syp/imp* double knockdown brains. This shows that *pros* is regulated directly by Syp, not via its downregulation of Imp. RNAi constructs are driven by insc-GAL4 (Betschinger *et al.,* 2006).

**Figure S2:**
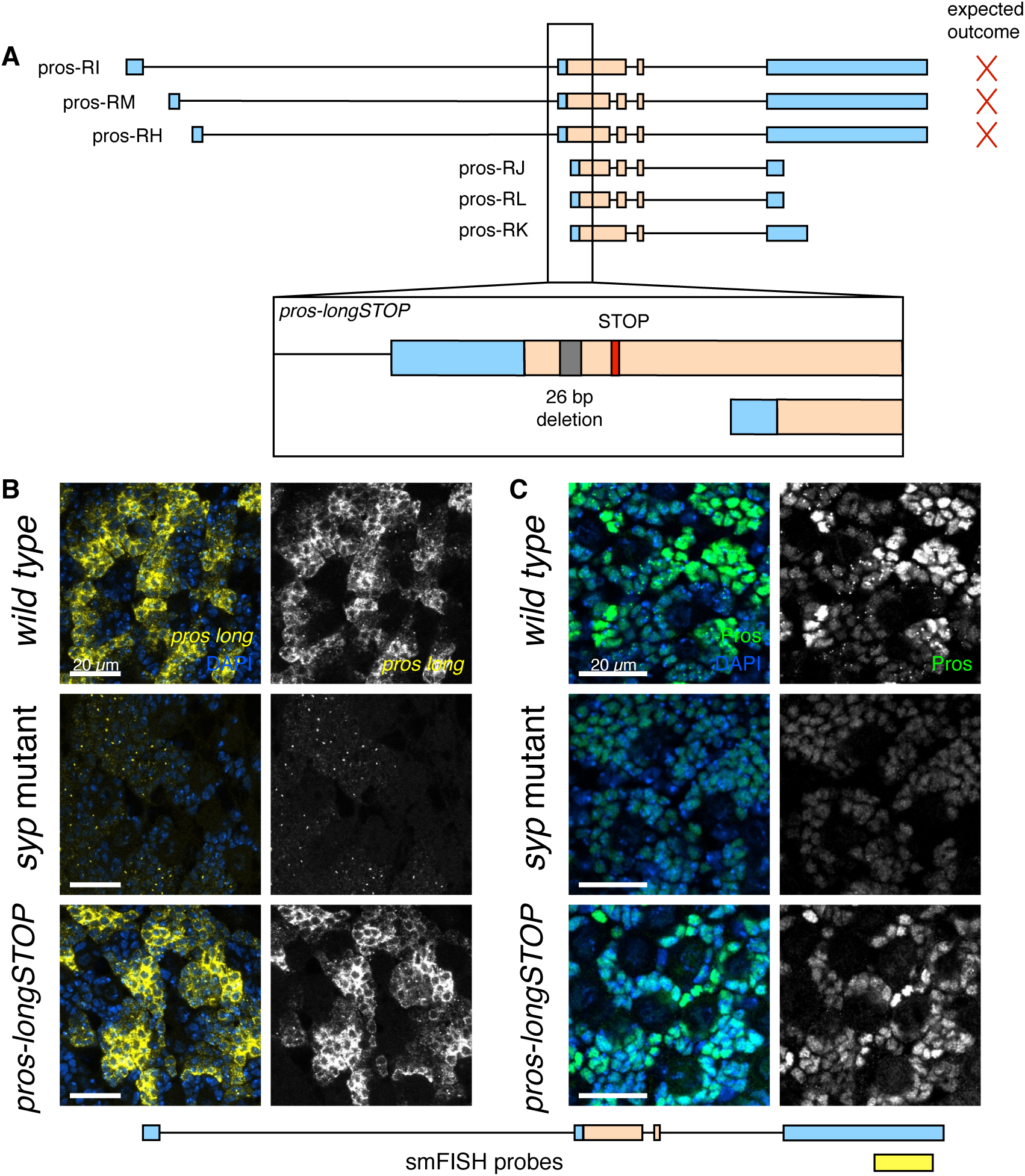
Introducing an NMD mutation does not remove *pros^long^*. **A** We used CRISPR/Cas9 to make a double stranded DNA break in the upstream coding region unique to the *pros^long^* isoforms. Repair by non-homologous end joining resulted in a 26 bp deletion, leading to a frame shift which introduces a premature stop codon 80 bp downstream (*pros-longSTOP*). This should produce a truncated polypeptide and induce NMD of the *pros^long^* mRNA isoforms. The *pros-longSTOP* mutant was stained with **B** *pros-long* smFISH and **C** Pros protein. Neither *pros^long^* or Pros protein were decreased compared to *wild type*.

**Figure S3:**
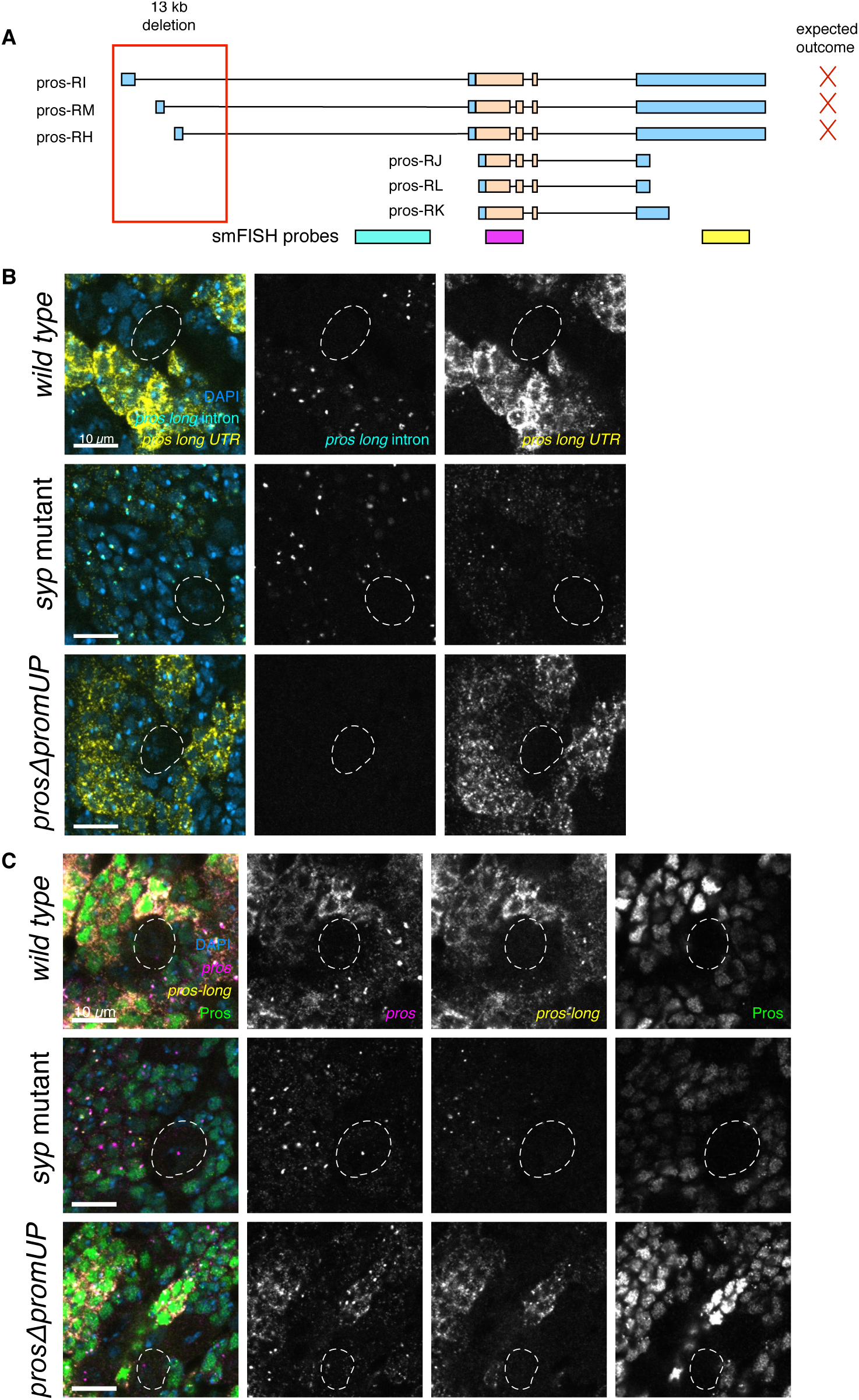
The deletion of the upstream *pros* promoters uncovers a new isoform. **A** We deleted the three upstream transcription start sites, which are annotated to produce *pros^long^.* (*prosΔprosUP*). **B** Staining the with *pros-long* intron and *pros-long* UTR smFISH showed that transcription from the upstream promoters was abolished, but low levels of *pros-long* UTR signal remained. **C** Staining with Pros protein additionally showed that the neuronal upregulation of Pros protein was not disrupted.

**Figure S4:**
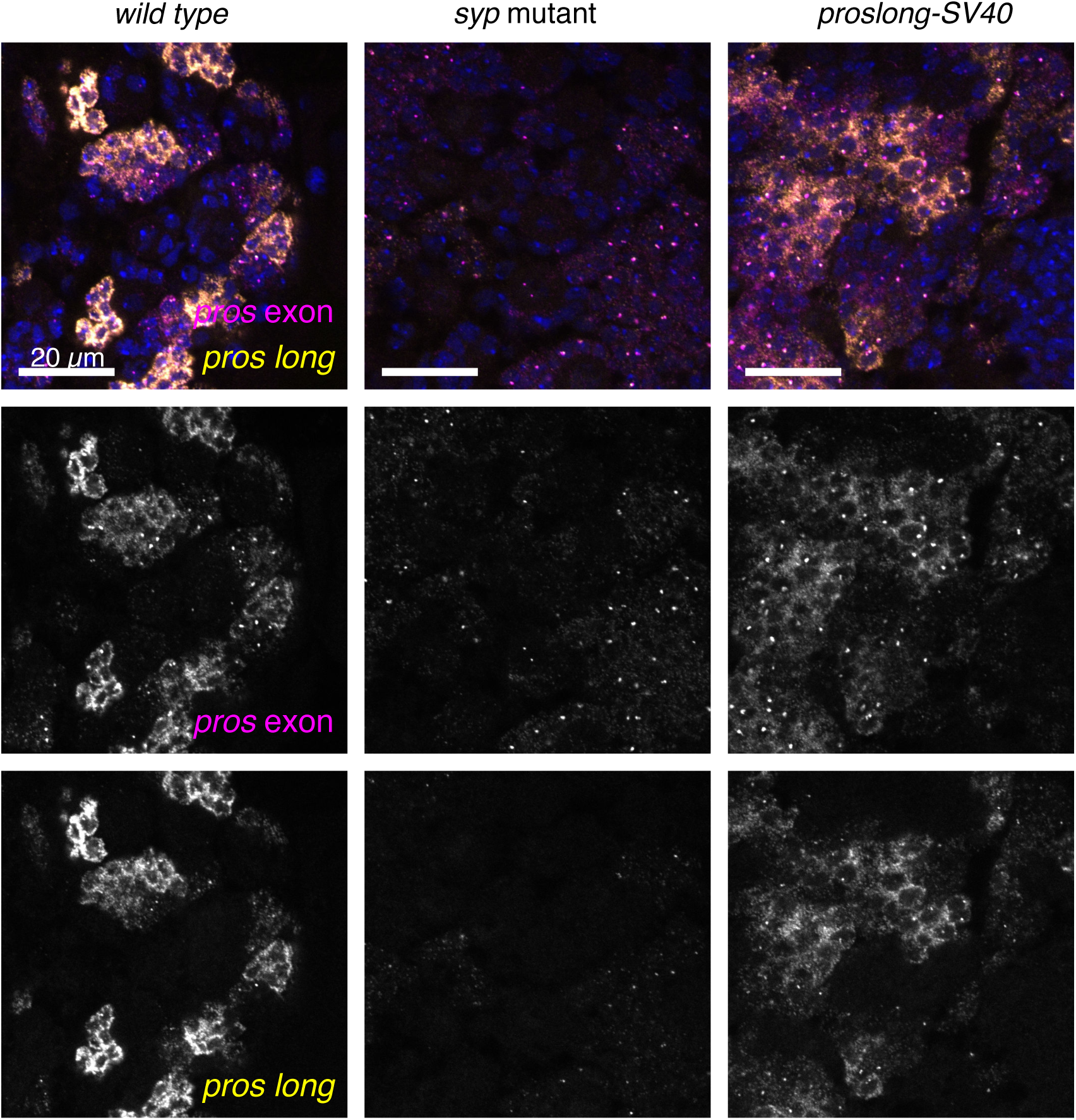
Insertion of an SV40 transcriptional terminator reduces, but does not abolish, *pros^long^* expression. Staining for *pros* exon and *pros-long* smFISH shows a reduction in *pros^long^* expression in the *proslong-SV40* line compared to *wild type*. However the *pros^long^* level is not reduced as much as in the *syp* mutant.

**Movie S1 - *proslong-REDr* flies exhibit defects in their activity**

1-3 day old mixed sex flies were transferred to empty vials and allowed to acclimatise for two hours without disturbance. The vial on the left contains *wild type* flies and the vial on the right contains *proslong-REDr* homozygous flies. After disturbance, by tapping the vial to bring all flies to the bottom, the *wild type* flies immediately crawl up the sides of the vial. However the activity of the *proslong-REDr* flies is impaired and the flies remain at the bottom of the vial.

**Table S1:**
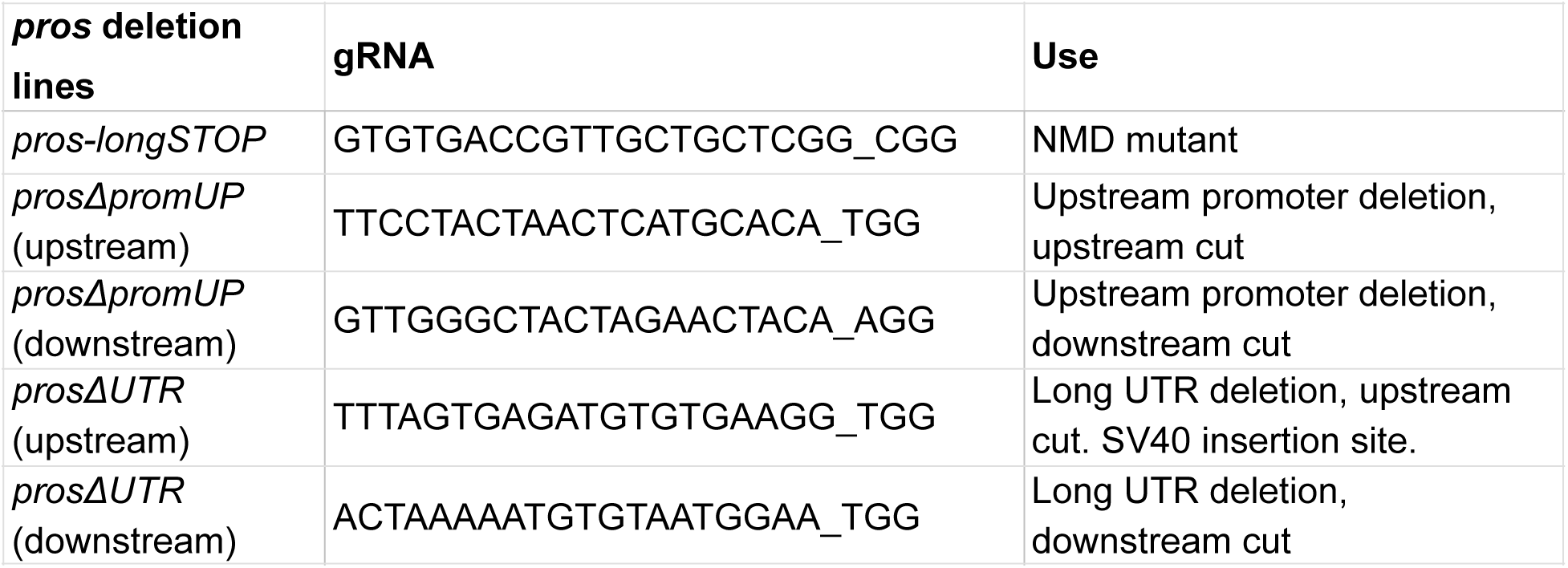
gRNA constructs. Guide RNA constructs used to produce the *pros^long^* deletion lines.

**Table S2:**
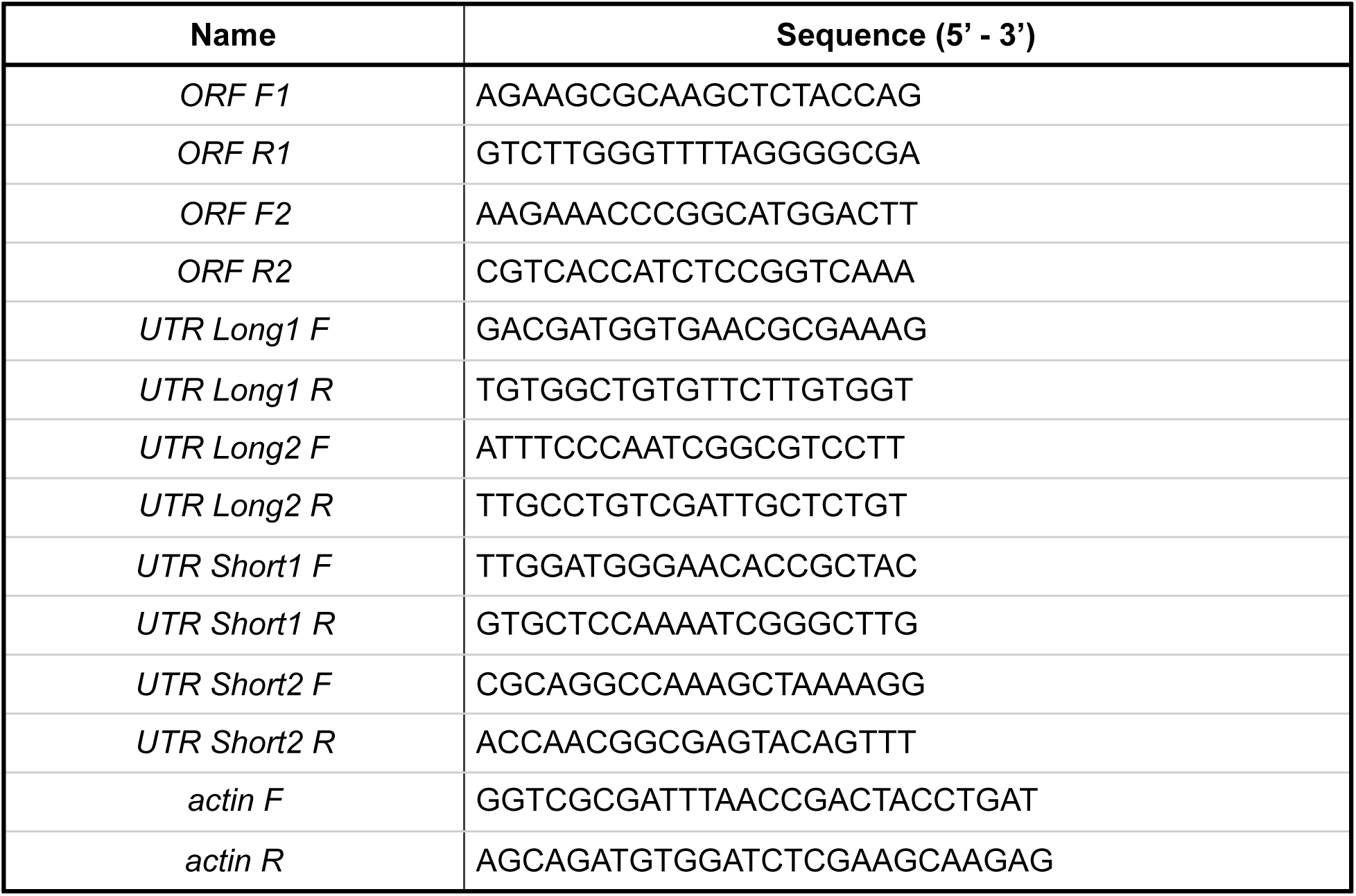
Primers for generating Northern blot probes. The above primers are used to generate radioactive Northern blot probes. ORF - Open Reading Frame of *pros* transcript; UTR Long are used to generate probes that are specific to the 15 kb *pros* 3’ UTR (probe LONG); UTR Short are used to generate probe that is against region that is common to all *pros* 3’ UTRs (probe ALL).

**Table S3:**
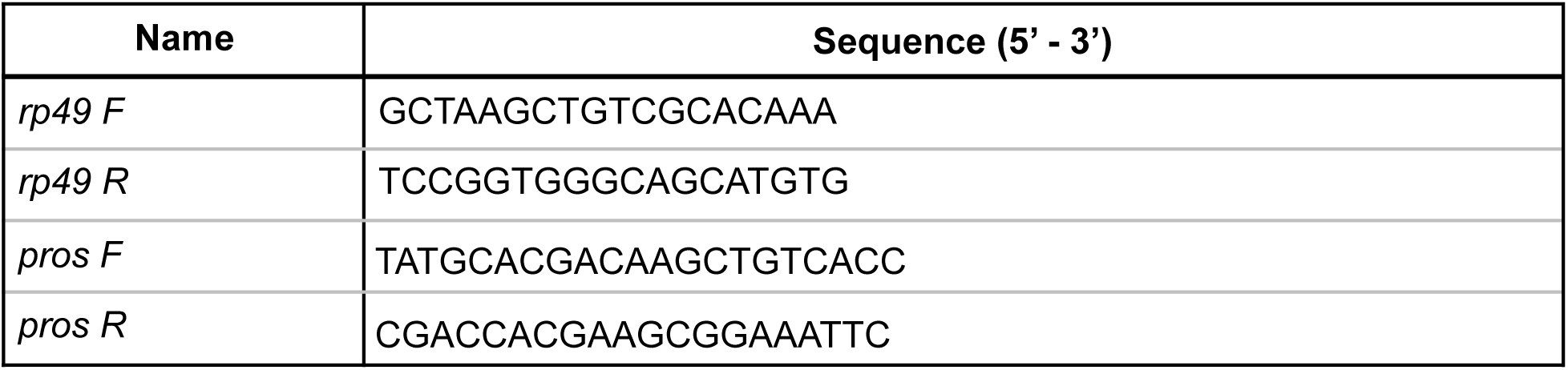
Primers used for RT-qPCR for Syncrip and IgG immuno-precipitation experiments. *pros* and housekeeping gene *rp49* was used to assess the efficiency and specificity of Syncrip binding to *pros*. Isoform specific primers and anchors were used for isoform specific RT-qPCR.

## METHODS

### KEY RESOURCES TABLE

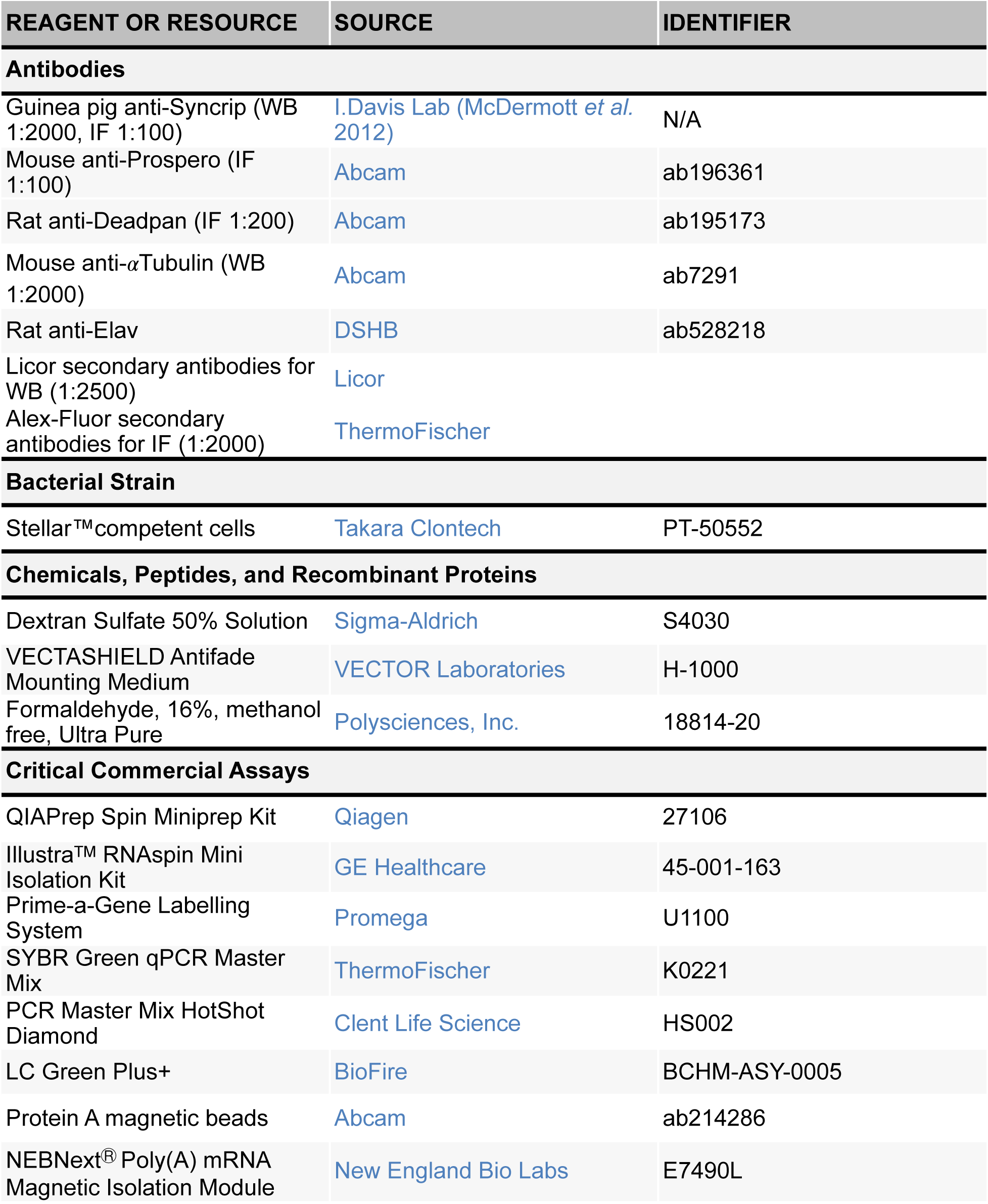
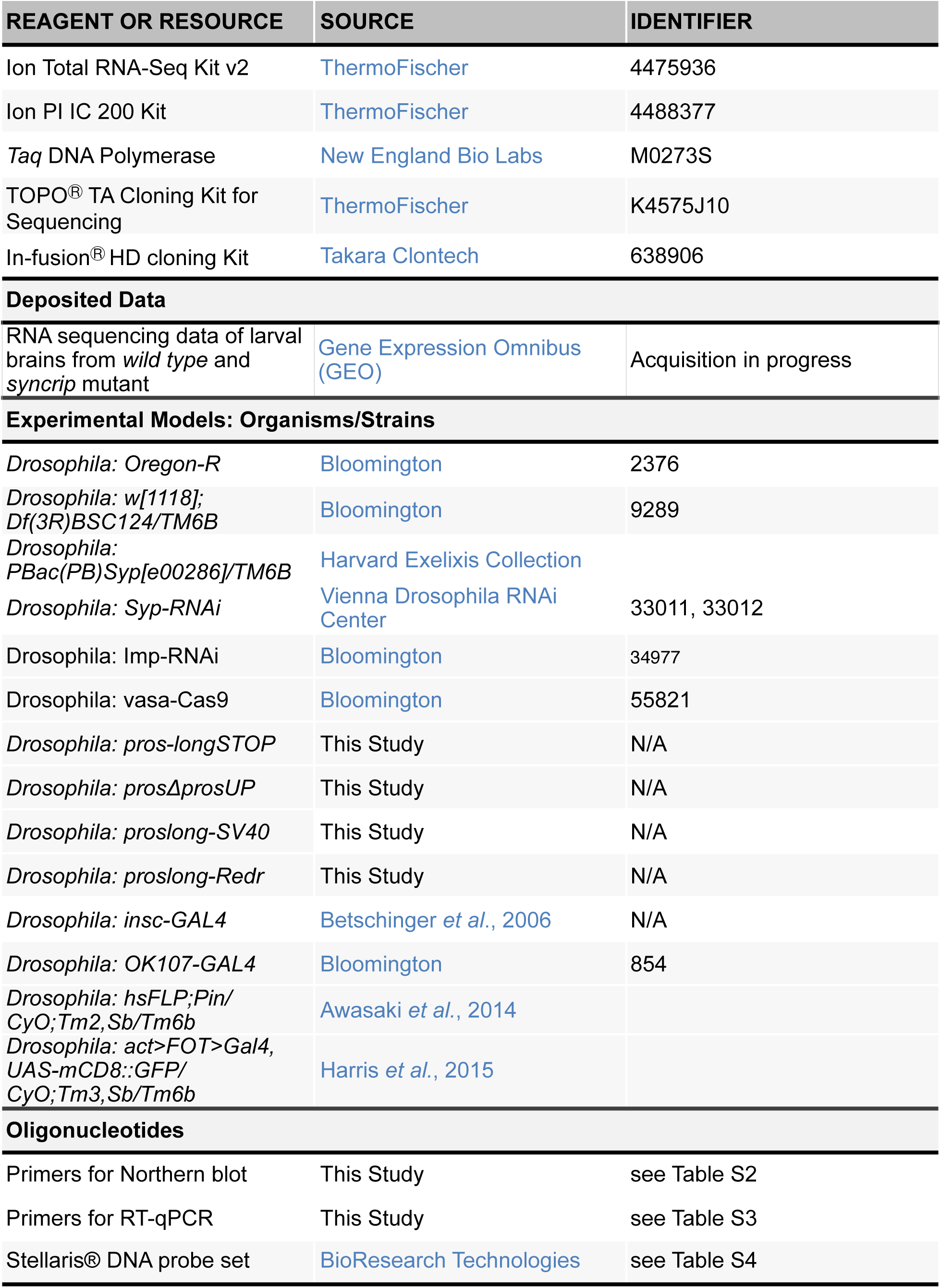
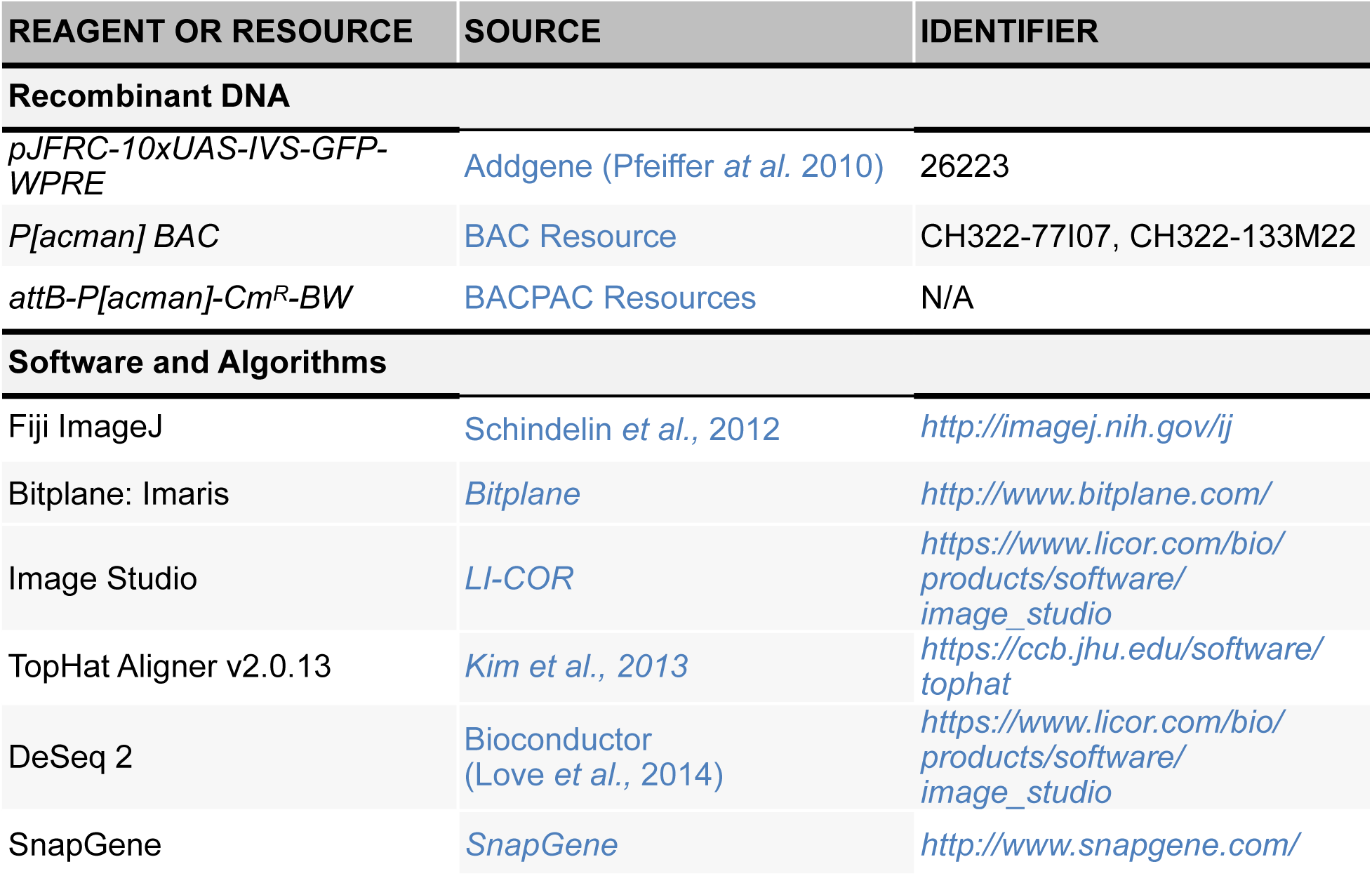

### CONTACT FOR REAGENT AND RESOURCE SHARING

For further information and access to any reagents and fly strains used in this manuscript, please contact corresponding author Ilan Davis.

### EXPERIMENTAL MODEL AND SUBJECT DETAILS

#### Fly Genetics

All strains were raised on standard cornmeal-agar medium at 25 ℃. Oregon-R (*OrR*) was used as the *wild type* strain.

#### Clonal analysis

For *wild type*, female hsFLP;Pin/CyO;Tm2,Sb/Tm6B were crossed to male act>FOT>Gal4, UAS-mCD8::GFP/CyO;Tm3,Sb/Tm6b. These lines were crossed to the syp mutant to produce parental lines to make the clones: female hsFLP;Pin/CyO;Syp[e00286]/Tm6b were crossed to male act>FOT>Gal4,UAS-mCD8::GFP/ CyO;Syp[e00286]/Tm6b. Eggs were collected in a four hour egg lay on apple juice plates with yeast paste at 25 °C. 24 hours after the egg lay, newly hatched larvae were transferred to fly food vials and incubated in a 37 °C water bath for 25 minutes. Wandering larvae were dissected 96 hours after larval hatching. For *syp* mutant, only non-tubby larvae were dissected. Immunofluorescence was completed as described below and clones were imaged on the ventral side.

#### *pros^long^* deletion mutant generation using CRISPR

sgRNA construct design and validation was performed by Dr. Andrew Bassett - Genome Engineering Oxford (GEO) (Table S1). Plasmids were injected into *vasa-cas9* embryos (Bloomington Stock: BL55821) as previously described (Bassett and Liu, 2014). For homologous recombination repair, the donor construct included a ds-RED marker for selection, flanked by lox sites to remove the dsRED, and an attP site to allow future insertions at the site. This inserted cassette was flanked by 1 kb homology arms to facilitate repair. The pDsRED-attP vector 51019 was used. Homology arms were amplified from purified BAC DNA (CH322-133M22).

##### 1. Nonsense Mediated mRNA Decay mutant

We took advantage of the unique coding sequence that is specific to the three long isoforms of *pros*, contributing an additional 300 amino acids to the N-terminus of Pros protein. We used CRISPR/Cas9 with non-homologous end joining to induce a frame shift which introduces a stop codon 80 bp downstream (Figure S2A). This stop codon would be expected to produce a truncated protein and induce Nonsense mediated mRNA decay (NMD) of the *pros^long^* transcripts. However this line (*pros-longSTOP*) showed no reduction in either *pros^long^*-specific smFISH signal (Figure S2B), or Pros protein (Figure S2C). This result suggests that the upstream coding sequence is not actually translated in *pros^long^* in the neurons. Instead, Pros protein is most likely produced from the downstream translation start site.

##### 2. UTR extension deletion

We aimed to directly delete the *pros* UTR extension using a CRISPR strategy with homologous recombination (HR), but this was unsuccessful. We speculate that the mutant line was embryonic dominant lethal, implying unknown roles for the UTR extension or non-coding transcription from this region.

##### 3. Upstream promoter deletion

We used an HR strategy to delete the three promoters which are annotated to produce the UTR extended isoform, collectively termed *pros^long^* (Figure S3A). We found that this line (*prosΔpromUP*) successfully abolished transcription from the upstream promoters, but the *pros^long^*-specific UTR probe showed significant remaining expression, albeit reduced compared to *wild* type (Figure S3B). Furthermore, the level of Pros protein was unchanged compared to *wild type* (Figure S3C). We conclude that the UTR extension of *pros^long^* can be transcribed from the downstream promoters. The lower levels of *pros^long^* in this line are sufficient to produce the *wild type* upregulation of Pros protein in neurons.

##### 4. Transcriptional termination

We designed a strategy to terminate transcription at the beginning of the 3’ UTR extension using a strong SV40 transcriptional terminator. This strategy uses a two-step process to insert an attP site and then integrate the SV40 terminator. The SV40 terminator (*proslong-SV40*) did not prevent the formation of the extended UTR of *pros^long^* (Figure S4). Unexpectedly, the inserted *dsRED* marker used to screen for integration does terminate the UTR extension of *pros* (Figure 6). We tested lines with the dsRED marker inserted in both orientations, which had the same phenotype. The *dsRED* inserted in the opposite orientation to *pros* (*proslong-REDr*) gave the lowest background and so was used for further experiments.

## METHOD DETAILS

### RNA extraction

Third instar larval brains were homogenised in IP buffer (50 mM Tris-HCl pH 8.0, 150 mM NaCl, 0.5% NP-40, 10% glycerol, 1 mini tablet of Complete EDTA-free protease inhibitor and 2 µl RNAse inhibitor (RNAsin Plus RNase Inhibitor, Promega). RNA extraction was performed using the RNAspin Mini kit (GE Healthcare).

### Western blot

NuPage Novex 4-12% Bis-Tris gels (Invitrogen) were used for all SDS-PAGE experiments. Following SDS-PAGE, proteins were transferred to nitrocellulose membrane using the XCell II™ Blot Module (Invitrogen) following manufacturer’s protocol. Protein bands were visualised with a quantitative infrared imaging system (LI-COR Odyssey).

### Northern blot

5 µl of purified RNA samples was mixed with 3.5 µl 37% formaldehyde and 10 µl 100% formamide, 2 µl supplemented 10x MOPS buffer (400 mM MOPS buffer, 100 mM NaOAc, 10 mM Na_2_EDTA). Sample mixture was incubated at 55 °C for 15 min and 4 µl of loading dye (1 mM EDTA, 0.25% BromoPhenolBlue, 50% Glycerol) was added and ran on an 1.5% agarose gel at 120 V in 1x MOPS buffer. ssRNA ladder (NEB) was used as size marker for all Northern experiments.

RNA was transferred to pre-wet Nylon membrane (Hybond-N, Amersham). Blotted RNA was cross linked to the membrane with either a UV crosslinker or by baking at 80 °C for >1 hr. The membrane was hybridised in hybridisation buffer (0.4 M Na_2_HPO_4_, 6% SDS and 1 mM EDTA) for 1 hr at 57 °C with RNA probes prepared using the Prime-a-Gene kit (Promega) (For list of primers used to generate RNA probes, see Table S2), washed twice with wash buffer 1 (40 mM Na_2_HPO_4_, 5% SDS and 1 mM EDTA) and twice with with wash buffer 2 (40 mM Na_2_HPO_4_, 5% SDS and 1 mM EDTA). RNA was detected overnight in phosphorImager (Molecular Dynamics).

### Real Time Quantitative PCR (RT-qPCR)

RT-qPCR was performed using a real time PCR detection system (CFX96 Touch^TM^ Real-Time PCR Detection System (BioRad)) and in 25 µl consisted of : 12.5 µl 2x SYBRGreen Mastermix (ThermoFisher), 0.75 µl of gene specific primer (10 µM stock, forward and reverse, Table S3), 4 µl of cDNA and 7 µl of nuclease free water. Cycle threshold (C(T)) value was calculated by the BioRad CFX software using a second differential maximum method. For each of three biological replicates, a dilution series of the input sample was produced (10%, 2%, 0.5%. 0.01% of input). qPCR for each set of primers was performed on this series and the Cq values were plotted against log10 dilution to find the formula of the line. The Cq value of each pulldown sample (anti-Syp and anti-IgG) was inputted into this formula to calculate the % of input pulled down. For each set of primers, a two tailed Student’s t-test was used to compare the % input pulldown in the test (anti-Syp) and control (anti-IgG) samples.

### Immunoprecipitation

Guinea pig anti-Syp antibody and IgG antibody were cross-linked to ProteinA magnetic beads (Abcam) following manufacturer’s protocol. For each replicate, 90 third instar larval brains dissected in Schneider medium were homogenised in immunoprecipitation (IP) buffer (50 mM Tris-HCl pH 8.0, 150 mM NaCl, 0.5% NP-40, 10% glycerol, 1 mini tablet of Complete EDTA-free protease inhibitor and 2 µl RNAse inhibitor (RNAsin® Plus RNase Inhibitor (Promega)) and topped up to 100 µl in IP buffer. For each reaction, 20 µl of lysate was taken as a 50% input sample which was taken directly to RNA extraction. 40 µl of lysate was incubated with 25 µl of each antibody cross-linked beads over night at 4 °C on rotator wheel. For each reaction, 100 µl of lysate was incubated with 20 µl of 50% bead slurry over night at 4 °C on rotator wheel. Next day, supernatant was transferred to fresh tubes and beads were washed 5 times for 5 min each with 200 µl cold IP buffer at 4 °C. After final wash, beads were resuspended in 40 µl extraction buffer (50 mM Tris-HCl pH 8.0, 10 mM EDTA and 1.3% SDS, 1:100 RNAsin) and incubated at 65 °C, 1000 rpm for 30 min on thermomixer. The elution step was repeated and the supernatant were pooled. RNA was then extracted from the IP eluates and the input sample and used for cDNA library synthesis.

### RNA Sequencing

Three biological replicates (*n* = 3), each replicate consists of a pool of 100 larval brains. Following RNA extraction, mRNA was enriched using NEB Next® Poly(A) mRNA Magnetic Isolation Module (NEB). Briefly, extracted RNA sample was mixed with Oligo d(T)_25_ beads and heated to 65 °C for 5 min followed by incubation at 4 °C for 1 hr to allow binding. Following incubation, the beads were washed 5 times for 5 min each at 4 °C and RNA was eluted by heating the beads at 80 °C for 5 min. Poly(A) enriched RNA was then used for library production using the Ion Total RNA-Seq Kit v2 for Whole Transcriptome Libraries (Life Technologies). Following quality control steps, adaptors were hybridised to the RNA fragments and RT reaction was performed followed by cDNA amplification with Ion Xpress RNA Barcode primers. Prior to sequencing, quality of cDNA libraries were assessed using Agilent High Sensitivity DNA Kit with the Agilent 2100 Bioanalyser. Libraries were pooled to a total concentration of 100 pM, with three samples multiplexed per chip. Sequencing was performed on an Ion Proton Sequencer, using the Ion PI IC 200 Kit (Life Technologies). Ion PI chips were prepared following manufacturer’s instructions and loaded using the Ion Chef System.

### Immunofluorescence

Third instar larval brains were dissected in Schneider’s medium and fixed in 4% formaldehyde (FA) solution (4% FA in 0.3% PBTX) for 25 min at room temperature. Following incubation with primary and secondary antibody, samples were mounted in VECTASHIELD anti-fade mounting medium (Vector Laboratories).

### RNA single molecule fluorescent *in situ* hybridisation (smFISH) for larval brains

Dissected 3rd instar larval brains were fixed in 4% paraformaldehyde (PFA) solution for 25 min at room temperature followed by 3 rinses in 0.3 % PBTX (PBS and 0.3% triton-X). Samples were washed 3 times for 15 minutes each in 0.3% PBTX at RT and incubated in pre-hybrdisation buffer (10% deionised formamide in prepared in 2x SSC) for 5 min at 37 °C. Hybridisation was performed by incubating samples overnight at 37 °C in the dark with gentle shaking in hybridisation buffer (10% deionised formamide, 5% dextran sulphate (sigma), 2x SSC) containing 250 nM gene-specific fluorescently labeled Stellaris® DNA probe set (BioSearch Technologies). Following hybridisation, samples were rinsed 3 times in pre-hybridisation buffer and washed for a further 3 times, 15 min each time in pre-hybridisation buffer at 37 °C. DAPI was included during the second 15 min wash. Samples were mounted in VECTASHIELD anti-fade mounting medium (Vector Laboratories) and imaged. Samples were protected from light for all steps including hybridisation.

### smFISH with immunofluorescence for larval brains

For smFISH with anti-Dpn antibody, the smFISH was performed identically as above, then after washing 3 times 30 minutes in prehybridisation buffer on Day 2, samples were blocked in blocking buffer (1% BSA in 0.3% PBTX) for 1 h at room temperature. Samples were incubated with primary antibody in blocking buffer for 3 h at room temperature, washed, and then incubated with secondary antibody for 1 h at room temperature. Final washes were performed using 0.3% PBTX, once including DAPI. For all other antibodies, the protocol is identical to the smFISH protocol with the following modifications. Before hybridisation, samples were blocked in blocking buffer (1% BSA in 0.3% PBTX) for 1 h at room temperature. Antibody at the appropriate dilution was included with the Stellaris® DNA probes during the over night hybridisation step. Counter-stain with secondary antibody was performed on the second day following hybridisation. After final wash, samples were mounted in VECTASHIELD anti-fade mounting medium (Vector Laboratories) and immediately imaged.

### Image acquisition

Larval brains were imaged on the ventral side. Fixed imaging of larval brain was performed using an inverted Olympus FV3000 Laser Scanning Microsope with Becker and Hickel FLIM system and with an inverted Olympus FV1200 Laser Scanning Microscope with high sensitivity galium arsenide phosphide (GaAsP) detectors (Olympus). Images were acquired using x20 0.75 NA UPlanSApo, x 40 1.3 NA Oil UPlan FLN, x60 1.4 NA and x100 1.4 NA Oil UPlanSApo objective lenses. Laser units used were: solid state 405 and 488 laser, argon 488, 515, 568 and 633 laser.

## QUANTIFICATION AND STATISTICAL ANALYSIS

### Bioinformatic analysis of RNA sequencing data

Base calling, read trimming and sample de-multiplexing was done using the standard Ion Torrent Suite. The reads were aligned to the *Drosophila* genome (ENSEMBL assembly BDGP5, downloaded 8 Jan 2015) using the TopHat aligner (v2.0.13) (Kim, 2013). To quantitate gene expression, uniquely aligned reads were assigned to the *Drosophila* exome (BDGP5) using htseq-count. Differential was assessed using negative binomial generalised linear models implemented in R/Bioconductor package DESeq2 (Love et al., 2014).

### Image analysis for smFISH

*pros* intron probes were used to compare *pros* transcription rates in *wild type* and *syp* mutant brains (Figure 3E). Transcription rates were compared separately for type I NBs and for progeny cells (including GMCs, immature and mature neurons). *pros* intron transcription foci were identified in 3D with the spot detection feature of Imaris Image Analysis software (*Imaris, Bitplane*), using a 1.2 µm diameter spot. Spots overlapping the edge of the image were removed. Average intensity was measured for each spot. In type I NBs, *pros* transcription foci are found spatially separated in the nucleus and therefore the intensity of two spots were added together for each NB. In progeny cells, the transcription foci are close in the nucleus and are measured in a single spot. Intensity measurements were normalised to the average *wild type* intensity for each experimental replicated (i.e. the average of *wild type* measurements from a single day of staining/imaging).

## DATA AND SOFTWARE AVAILABILITY

We are currently acquiring a Gene Expression Omnibus (GEO) accession number for the presented RNA sequencing data and will provide the number prior to publication.

